# A natural transdifferentiation event involving mitosis is empowered by integrating signaling inputs with conserved plasticity factors

**DOI:** 10.1101/2021.05.05.442643

**Authors:** Claudia Riva, Martina Hajduskova, Christelle Gally, Arnaud Ahier, Sophie Jarriault

## Abstract

Transdifferentiation, or direct cell reprogramming, is the direct conversion of one fully differentiated cell type into another. Whether core mechanisms are shared between different transdifferentiation events, which can occur naturally in presence or in absence of cell division, is unclear. Our lab has previously characterized the Y-to-PDA natural transdifferentiation in *Caenorhabditis elegans*, which occurs without cell division and requires orthologs of vertebrates’ reprogramming factors. In this study, focusing on another transdifferentiation process, the K rectal cell-to-DVB GABAergic neuron, we report that the Y-to-PDA reprogramming factor SEM-4/SALL, SOX-2, CEH-6/POU are required for K-to-DVB transdifferentiation to allow the erasure of the rectal identity. In addition, cell division is necessary but not sufficient for this transdifferentiation event while the Wnt signaling plays distinct functions during the process including the selection of the daughter cell with a different fate, loss of the rectal identity and imposition of the specific neuronal subtype identity. We provide evidence that both the Wnt signaling and Y-to-PDA reprogramming factor SEM-4/SALL, SOX-2, CEH-6/POU act in parallel for the rectal identity erasure. Our results further support a model where antagonistic activities of SOX-2 and POP-1 and decreasing SOX-2 levels over time provide a timer for the acquisition of the final identity. In addition, the different levels of SOX-2 provide a mechanism for the integration of Wnt opposite dedifferentiation and re-differentiation functions during K-to-DVB transdifferentiation.

## INTRODUCTION

The maintenance of stable cell identities is fundamental to ensure the function of tissues and organs, and to avoid oncogenic transformation (Roy and Hebrok, 2015). Nevertheless, cell identity changes have been observed, for instance during regeneration after an injury to replace cell types that have been lost (Brockes and Kumar, 2002; Thorel et al., 2010; Yanger et al., 2013). Cell identity conversion of differentiated cells have been described: one such event is transdifferentiation (Td, also known as direct cell reprogramming), the process through which a differentiated cell is reprogrammed directly into another differentiated cell (Eguchi and Kodama, 1993). This phenomenon has been described across phyla under physiological and pathological conditions (Jessen et al., 2015) and together with observations as early as the eighteen century on the flexibility of differentiated identities (Trembley 1994, Virchow, 1886), has challenged the idea that terminal differentiation can be irreversible (Merrell and Stanger, 2016). The final blow to the notion of irreversibility of terminal cell identity was later provided by Takahashi and Yamanaka, 2006 who showed that cocktails of pluripotency transcription factors (TFs) could reprogram somatic cells into pluripotent stem cells *in vitro*.

Induced reprogramming of somatic cells *in vitro* into other somatic cell types can also be achieved but is often inefficient and difficult to track at the single cell level, leaving the lineal relationship between the initial and final cell identities ambiguous. Natural Td, on the other hand, is robust in terms of reprogramming efficiency (Eguchi et al., 2011) and offers the opportunity to decipher the cellular and molecular mechanisms at play in a complex tissue, where cell autonomous and non-autonomous signals are involved. Our lab has been taking advantage of *C. elegans* to study natural Td *in vivo* at the single-cell resolution. *C. elegans* transparency and stereotyped cell lineage (Sulston and Horvitz, 1977) offers a powerful tool to identify and unambiguously follow cell plasticity phenomena. We have previously described and characterized a natural Td event occurring in 100% of the animals during *C. elegans* larval development, where the Y rectal cell becomes a motor neuron called PDA without cell division nor cell fusion (Jarriault et al., 2008). This event occurs in a stepwise process that starts with the erasure of the initial identity, before re-differentiating into a different cell type (Richard et al., 2011). Through unbiased EMS and RNAi screens, we have identified sets of genes that mediate and tightly control the different steps of the process (Kagias et al., 2012; Richard et al., 2011; Zuryn et al., 2014). Interestingly, orthologs of some factors involved in Y dedifferentiation, e.g. CEH-6/POU, SOX-2/SOX and SEM-4/SALL (Jarriault et al., 2008; Kagias et al., 2012), are known to have reprogramming activities or to be associated to pluripotency in mammalian cells (Julian et al., 2017; Malik et al., 2018; Ng and Surani, 2011). This led us to ask whether these factors could constitute a conserved plasticity cassette that could be shared with other Td events. Furthermore, even though Y-to-PDA does not require cell division, both natural and induced Tds often involve mitosis (Lambert et al., 2021). Whether the mechanisms required for Td in presence or absence of cell division remain the same is still an open question.

In this study, we aimed at addressing these questions by characterizing and dissecting another putative Td event in the worm rectum, that occurs in the presence of a cell division. This event involves the K rectal cell, which is born during embryogenesis and divides during the L1 stage. K division gives rise to an anterior daughter, K.a, which remains part of the rectum and to the posterior daughter K.p that becomes in the early L2 stage a GABAergic neuron called DVB (Fig. 1A). Using *in vivo* imaging and electron microscopy we characterized K, K.a, K.p and DVB identities and confirmed that K-to-DVB is a *bona fide* Td event. We showed that K division is oriented and asymmetric and is crucial for DVB formation. Using gene candidate approaches and cell identity markers, we demonstrated that it involves the *C. elegans* Wnt/β-catenin asymmetry pathway. The exterior WNT signal ensures that it is the K.p daughter, formed at a stereotyped posterior position, that loses its mother’s identity. However, the Wnt signaling pathway alone is not sufficient to drive K-to-DVB. We found that the plasticity factors required for Y-to-PDA Td (Kagias et al., 2012) are also required for K-to-DVB Td for and after the division. By dissecting the genetic relationships between plasticity factors and the Wnt signaling pathway, we found that Wnt most likely acts in parallel to SEM-4 to erase the initial rectal epithelial identity of K.p. Finally, through the analysis of POP-1/TCF and SOX-2 binding sites in *lim-6* regulatory sequences, our data suggest that Wnt and SOX-2 provide a developmental timer to refine the re-differentiation step through direct regulation of the expression of the DVB terminal selector *lim-6*. Thus, our study provides an integrated view of how a fully differentiated cell can be reprogrammed into a totally different cell type at the single cell resolution and shows the existence of core “plasticity factors” required for Td independently of the presence or absence of cell division. Moreover, it shows how context-dependent signaling pathways may contribute to the cellular environment in parallel to this plasticity cassette, to lead to different dynamics and different final cell identities.

**Fig. 1.**
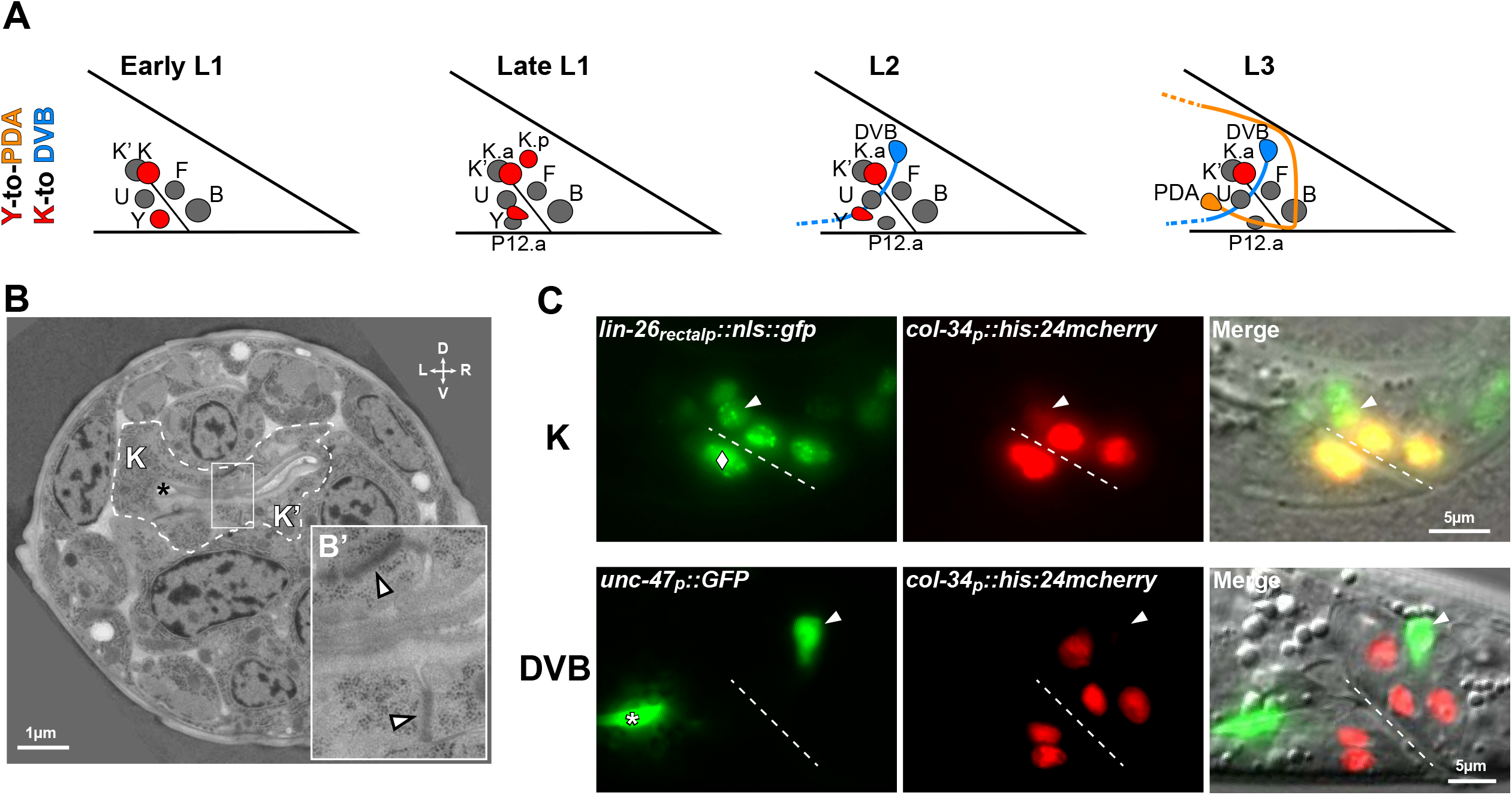
The K-to-DVB cellular process is a new transdifferentiation model. **(A)** Schematic illustration of the rectal area during the formation of PDA and DVB neurons from the Y and K rectal cells resp. in hermaphrodites. In early L1, both Y and K are part of the rectum, a tube made of three bi-cellular rings (Y-B, U-F, K-K’). In late L1, K divides into two daughter cells: K.a and K.p. In L2, Y starts to retract from the rectum while P12.pa replaces it in the rectum; K.a is part of the rectum in place of its mother and K.p becomes DVB. In L3, PDA is formed. Rectal cell nuclei are represented; anterior is to the left and dorsal up. **(B)** Electron micrograph of the rectal area of a newly hatched L1 hermaphrodite showing the K and K’ cells outlined with the dashed white line. Black star, rectal lumen; white arrowheads, electron-dense apical junctions. (B’) Magnification of the boxed area in B, illustrating an electron-dense apical junction between K and K’. D, V, L, R: dorsal, ventral, left and right orientations. **(C)** Fluorescent and DIC images of nuclear *lin-26p::GFP* in the rectum of an L1 animal (top) and *unc-47p::GFP* in GABAergic neurons in an L3 animal (bottom). *col-34p::his-24::mcherry* allows the visualization of the rectal cells nuclei and K.p after its birth. The white diamond in L1, migrating Y rectal cell partially overlapping U; white star in L3, VD13 GABAergic neuron; arrowheads, K in L1 and DVB in L3; dashed line, rectal slit. Anterior is to the left and dorsal up.

## RESULTS

### K-to-DVB cell identity conversion involves a differentiated rectal cell that gives rise to a neuron

Like for Y becoming the PDA neuron (Jarriault et al., 2008), the determination of the embryonic and post-embryonic somatic cell lineage in *C. elegans* (Sulston et al., 1983) has allowed us to pinpoint putative cell fate changes occurring during *C. elegans* larval development. Here we focus on the K rectal cell which gives rise to two daughter cells through a single cell division in late L1 stage (Fig. 1A): K.a, which replaces K in the rectum, and K.p, which subsequently becomes a GABAergic motor neuron called DVB (McIntire et al., 1993). We first characterized in depth the identities of the initial K mother cell and the final DVB cell in hermaphrodites by analyzing K and DVB cellular morphologies and the expression of specific cell-identity marker genes.

K is born in the embryo and forms one of the three rectal rings with its sister K’ through adherens junctions (Sulston et al., 1983). The six *C. elegans* rectal cells are differentiated, specialized epithelial cells which form the *C. elegans* rectum (<u>wormatlas</u>). The K rectal identity is confirmed at the ultra-structural level in early L1 larvae, where K shows a typical rectal-epithelial morphology resembling the one of K’ (Fig. 1B). Moreover, it expresses several epithelial and rectal markers, while it does not express any neuronal marker (Fig. 1C, Table S1, Ferreira et al., 1999; Jarriault et al., 2008; Labouesse et al., 1996). Importantly, even though K is not post-mitotic and therefore is usually called a blast cell (Chisholm, 1991) it is not a stem cell: it is a mature rectal cell indistinguishable from the other rectal cells which do not divide nor change identity and with which it forms the rectum, a vital organ. Thus, while not post-mitotic, K is fully differentiated and holds a structural role in a permanent organ. On the contrary, DVB lacks epithelial and rectal markers, expresses neuronal genes, and develops in a typical neuronal morphology (White et al., 1986) to fulfil its GABAergic motor neuron function in defecation (Fig. 1C, Table S1).

Altogether, these observations demonstrate that K and DVB display different terminal identity features, the former being a fully differentiated rectal cell and the latter being a GABAergic neuron. The development of a neuron from a fully differentiated rectal cell is intriguing and reminiscent of the Y-to-PDA Td (Jarriault et al., 2008), and is in agreement with the original definition of Td by T. S. Okada (Okada, 1991). Thus, we investigated the mechanisms underlying DVB formation and the possible shared aspects with Y-to-PDA Td.

### K division, which is oriented and asymmetric, is necessary for DVB formation

Since K-to-DVB conversion, differently from Y-to-PDA Td (Jarriault et al., 2008), involves a cell division, we started by investigating the role of cell division in this process. First, we assessed whether K division *per se* is necessary for the formation of DVB. To this end, we used *lin-5/NuMA* mutant as we found that K cytokinesis is blocked in 90.5% of *lin-5/NuMA(ev571ts)* thermosensitive mutant animals raised at the restrictive temperature (25°C) (Fig. 2A-B) (Izumi et al., 2006). We observed that DVB never formed (Fig. 2B) in L4 *lin*-5 mutants in which K did not divide, while the worms retained a complete rectum. These results show that K division is necessary for the worms to develop DVB. Moreover, as already observed in other *C. elegans* cells (Lorson et al., 2000), we found that DNA replication in K occurred in most of the worms in which K cytokinesis failed, suggesting that this process is not enough for DVB formation from K in absence of cytokinesis (Fig. S1A).

**Fig. 2.**
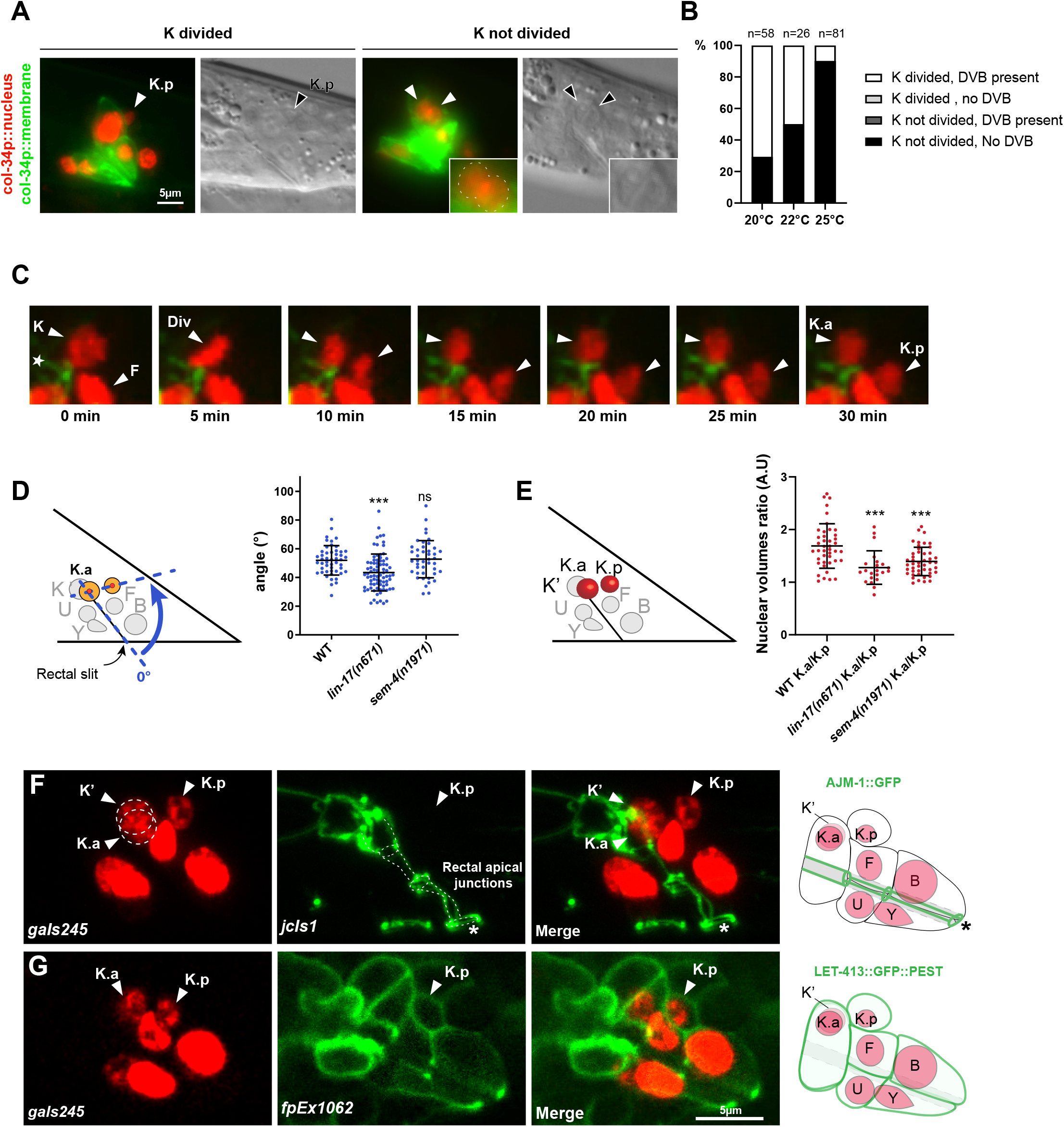
K division is oriented, asymmetric and gives rise to a K.p cell that retains epithelial features. **(A)** Fluorescent and DIC images of *col-34p::his-24::mcherry* (*col-34p::nucleus*) and *col-34p::ph::gfp* (*col-34p::membrane*) in *lin-5(ev571ts)* mutants where K division has (left) or has not (right) occurred. Absence of K cytokinesis in the worm on the right is evidenced by the presence of a unique cytoplasmic membrane around two nuclei. **(B)** Histograms summarizing both the occurrence of K division and later the percentage of absence of DVB in single *lin-5(ev571ts)* mutants at different restrictive temperatures, through a score-recover-score strategy. No orientation defects were observed in the few mutants where K divided. n, total number of animals scored. **(C)** Spinning time-lapse imaging of K division using *col-34p::his-24::mcherry* and *hmr-1::GFP* to visualize rectal cell nuclei and apical junctions resp. Time interval, 5min. Div, metaphase of K division; Arrowheads, K, K.a and K.p nuclei; white star, apical junction. (**D, E**) Quantification of the angle of K division with respect to the rectal slit (D), and Quantification of the nuclear volumes of K.a and K.p using *col-34p::his-24::mcherry* to visualize the nuclei (E) in late L1 wild type, *lin-17(n671)* and *sem-4(n1961)* animals. Bars represent mean +/-standard deviation. (D) Blue dashed lines (left), landmarks used for the measurements of the angles (curved arrow). 49, 79 and 43 animals were scored in wild type, *lin-17* and *sem-4* mutants, resp. (E) Dot plot representing the ratio for K.a/K.p in arbitrary units. 42, 22 and 46 nuclei were measured in wild type, *lin-17* and *sem-4* mutants, resp. **(F)** Confocal images representing the apical junction marker AJM-1::GFP (*jcIs1*) in a late L1 wild type larvae carrying *gaIs245[col-34p::his-24::mcherry]* to visualize rectal cell nuclei. Apical junctions (dashed lines) are present between the rectal cells and along the rectal lumen but not in K.p. Arrowheads, K, K.a and K.p nuclei; white star, rectal opening. **(G)** Confocal images representing the localization and expression of a destabilized basolateral marker LET-413::GFP::PEST (*fpEx1062*) in K.p in a late L1 wild type larvae. Rectal cell nuclei are visualized with *gaIs245[col-34p::his-24::mcherry]*. For all images, anterior is to the left, dorsal up.

Given the requirement of K cell division for DVB development, we next characterized it using time-lapse spinning disk microscopy. Using *gaIs245[col-34p::his-24::mcherry]* and *fpIs17[hmr-1::gfp]* transgenes to visualize the rectal cells’ nuclei and their apical junctions respectively and *oxIs12[unc-47p::gfp]* transgene to monitor DVB formation, we observed that K.p buds off from the K cell posteriorly, above the rectal cell F, without disrupting K apico-basal polarity (Fig. 2C, S2A). We did not observe any rounding of the K cell as often associated with cell division (Cadart et al., 2014), nor loss of adherence of K to its neighboring K’ cell: this division modality likely allows the maintenance of the integrity of the rectum during cell division. K division appears to always occur with an antero-posterior orientation; thus, we further analyzed this parameter in a quantitative manner. As K division is very fast, we estimated its orientation by measuring the angle formed by the rectal slit with the K.a and K.p’s nuclei alignment, maximum 1h after division. Our results show that orientation of K division is stereotyped among animals, forming an angle of 51.2±7.6° (Fig. 2D). Additionally, quantification of the nuclear volumes of K.a and K.p 1h after K division, using the *gaIs245*[*col-34p::his-24::mcherry*] chromatin marker as an approximation revealed an asymmetry in K.a and K.p nuclear volumes with K.a nucleus being 1.7-fold bigger than K.p nucleus (Fig. 2E). The stereotyped orientation and the adherence of K to its sister K’ during division led us to check whether apical junction proteins are asymmetrically segregated during K cytokinesis: indeed, as the orientation implied, we found that only the anterior daughter K.a, which takes on K function in the rectum, inherits the apically localized proteins AJM-1, HMR-1 and DLG-1, while they are absent in the posterior daughter K.p. Thus, K division appears oriented and asymmetric (Fig. 2F; Table S1).

This result prompted us to investigate whether a neuronal daughter is directly produced by cell division or whether K.p becomes neuronal later. To this end, we first determined the timeline of the cellular events occurring during K-to-DVB transition. We observed that K divides around 11.5h post hatching (PH) at 20°C. Around 4h later, after the L1-to-L2 molt which corresponds to 15-16h PH, we started to observe *unc-47* expression in the K.p daughter in a few young L2 larvae, and 16 to 17h PH we were able to detect *unc-47p::gfp* in all the L2s scored (Fig. S2A, B). Conversely, K, K.a and newly born K.p never express *unc-47p::gfp*. Since *unc-47*/*SLC32A1* is involved in GABA transport and necessary for GABAergic neuronal activity (McIntire et al., 1993), we will henceforth use “K.p” when considering K posterior daughter until 16h PH, and “DVB” after this time point. These observations suggest that K.p is not a differentiated GABAergic neuron at birth.

We further focused our attention on K.p identity. We observed that the basolateral epithelial marker LET-413 is present in K.p at the same level as in K and K.a in all the animals observed (Fig. S2B, Table S1). This presence is also visualized when a destabilized version of LET-413::GFP fusion protein is used, suggesting active expression of *let-413* gene (Fig. 2G). The proportion of animals with LET-413::GFP in K.p decreases to less than 20% after 16h PH when the DVB marker *unc-*47 is expressed in all the worms (Fig. S2B). On the same line, transcription of the epithelial TF *lin-26* is also observed in K.p after the division (Fig. S2B) and disappears quickly, since *gfp* expression is absent in about 20% of the worms from 1h after K division and onwards. Such GFP expression does likely not represent perdurance of the GFP protein since previous antibody staining data showed presence of LIN-26 in K.p (Labouesse et al., 1996). Moreover, we examined the presence of rectal markers in K.p and found that *egl-5/HOX, col-34*, and *got-1*.*2/GOT1* are present (Table S1). On the contrary, we observed that both pan-neuronal (like *unc-33/DPYS* and *unc-119/UNC119*) and GABAergic terminal differentiation genes (like *unc-25/GAD*) are not expressed in K.p after the division, but rather appear later (Table S1).

Altogether, these data show that K division is required to form the DVB neuron and is oriented with respect to the rectal tube, resulting in the asymmetric partitioning of apical junctions between the two daughter cells. Although K.p does not inherit apical junction proteins and shows a smaller nucleus than K.a, it retains important epithelial and rectal features, inheriting and expressing basolateral, rectal and hypodermal factors. Thus, successive steps are required to convert K.p into a GABAergic motor neuron.

### The Wnt/β-catenin asymmetry pathway is required for K-to-DVB

The stereotyped and oriented nature of K division prompted us to investigate the mechanisms regulating it and their impact on DVB formation. We focused our attention on the Wnt signaling pathway, known to orient the mitotic spindle (Schlesinger et al., 1999) and regulate several cell divisions in *C. elegans* resulting in asymmetric fates (Mizumoto and Sawa, 2007; Sawa and Korswagen, 2013). To test its potential role during K-to-DVB conversion, we first looked for DVB formation defects in mutant backgrounds affecting different components of the Wnt signaling pathway. *lin-17* encodes one of the four Frizzled receptors in *C. elegans* and is expressed in the rectal area (Sawa and Korswagen, 2013). We observed a strong defect in the severe loss-of-function *lin-17(n671)* mutant (Sawa et al., 1996), with more than 90% “NO DVB” worms (Fig. 3A). We next analyzed the requirement of several other Wnt pathway components: *lin-44* and *egl-20* which encode WNT ligands expressed in the rectal area (Harterink et al., 2011); *pop-*1 which encodes the unique T-cell factor/lymphoid enhancer factor (TCF/LEF) transcription factor in *C. elegans* (Lin et al., 1995); *bar-1, sys-1* and *wrm-1* which encode the three worm β-catenins involved in the Wnt signaling transduction downstream of Frizzled (Sawa and Korswagen, 2013). We found that *pop-1/TCF, lin-44*/*WNT, sys-1*/β-catenin and *wrm-1*/β-catenin are required for K-to-DVB to various degrees, while *egl-20/WNT* and *bar-1/*β-catenin are dispensable (Fig. 3A). Thus, the Wnt signaling pathway and, in particular components of the Wnt/β-catenin asymmetry pathway (Mizumoto and Sawa, 2007), are required to form the DVB neuron.

**Fig. 3.**
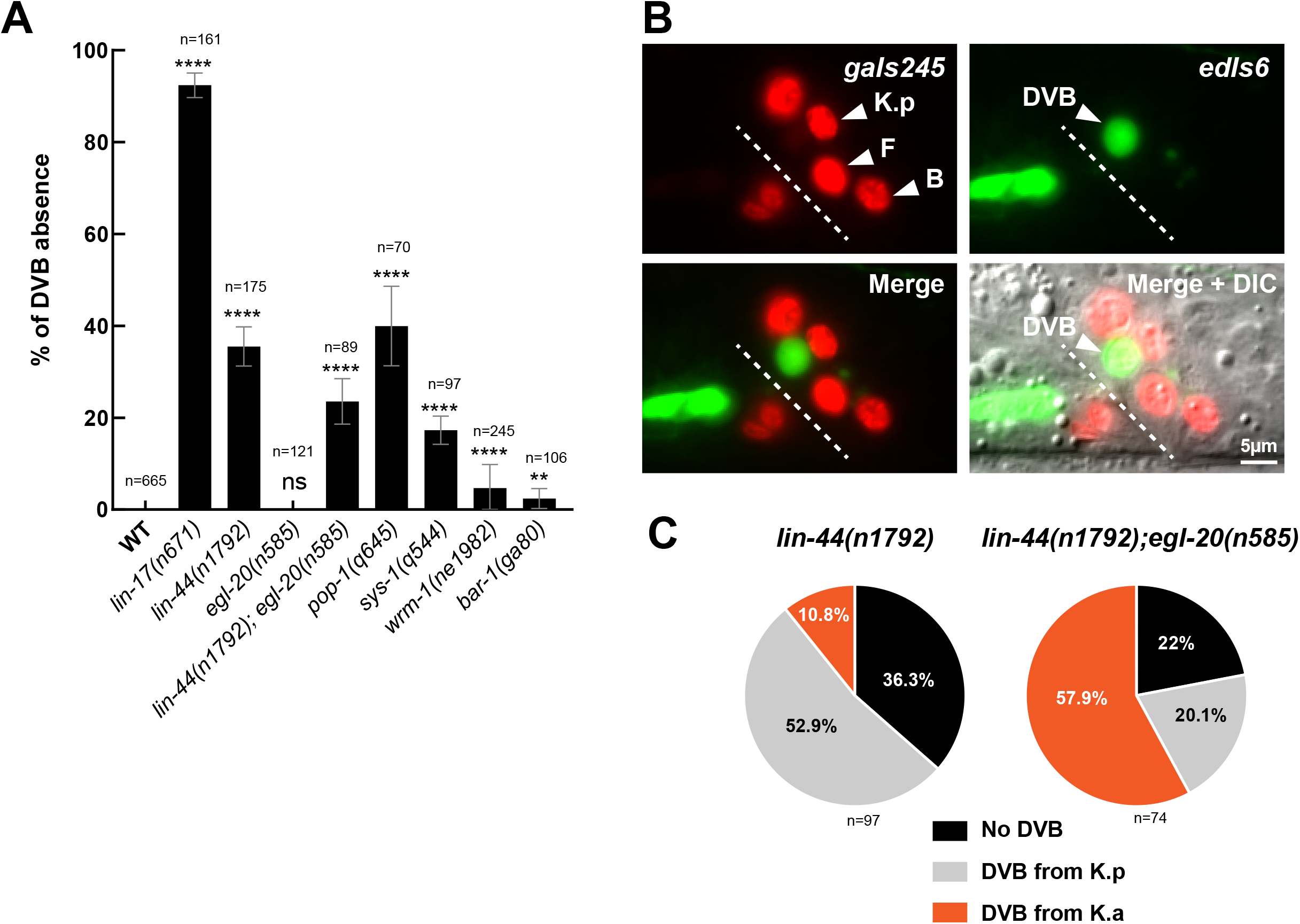
The Wnt signaling is involved in DVB formation. **(A)** Mutations in Wnt signaling pathway components lead to DVB defects. The frizzled receptor *lin-17* mutant shows the strongest DVB formation defect, while the β-catenin *bar-1* or the Wnt *elg-20* mutants display low or no DVB defect. n, total L4 larvae scored. **(B)** Fluorescence microscopy pictures of a double mutant *lin-44(n1792); egl-20(n585)* L4 worm where DVB appears to be formed from K anterior daughter. *gaIs245* highlights the rectal cells’ nuclei and *oxIs12*, the GABAergic neurons including DVB. **(C)** Representation of the percentages of the indicated phenotypes in *lin-44(n1792)* single and *lin-44(n1792); egl-20(n585)* double mutants shown in (A). When formed, DVB is mostly formed by K.a in the double mutant strain. n, total animals scored.

We next examined if K division was altered in worms with defective Wnt signaling. We found that division occurred in all *lin-17(n671)* mutant animals observed (n=103). Thus, the highly penetrant absence of DVB seen in this mutant is not due to a failure in K division. However, we found that the orientation of K division is affected in 8.9% of the *lin-17/FZD* mutant animals (Fig. 2D), with K.p positioned more dorsally or ventrally. Although not accounting for the total “NO DVB” defects in *lin-17/FZD* mutants, these data suggest that K division is abnormal in a fraction of them. We further tested the involvement of other Wnt-related pathways, such as the non-canonical Wnt pathway, known to act directly on spindle orientation through *ced-10/RAC1* (Cabello et al., 2010; Schlesinger et al., 1999), and of *lin-18/RYK, cam-1/ROR* and the Planar Cell Polarity (PCP) pathway (*vang-1/VANGL* and *fmi-1/CELSR2*) (Sawa and Korswagen, 2013). We found that none of these genes is involved in K-to-DVB (Fig. S3A, B).

To directly test whether the orientation of K division has an impact – at least partially – on DVB formation, we aimed at perturbing it using mutants known to randomize the cell division axis (Gotta and Ahringer, 2001) and analyzing the impact on K-to-DVB conversion. Since all temperature conditions tested for the *lin-5(ev571ts)* mutant resulted in either wild type DVB or “NO K cytokinesis/NO DVB”, precluding its use for our purpose (Fig. 2B), and *gpr-1* and *gpa-16/GNAI1* mutants showed no defect at all (Fig. S1B), we examined the division angle in the 0.9% *goa-1(sa734)* null mutant displaying “NO DVB” defect (Fig. S1B). Using a score-recover-score approach, we found that 8.6% of the animals exhibited an abnormal K division angle at late L1 stage (Fig. S1C) that did not translate into any “NO DVB” defect. Thus, while LIN-17/FZD activity may impact on K division axis, altered orientation of K division in itself does not seem to affect DVB formation.

Altogether, these results show an involvement of the sole canonical Wnt/β-catenin asymmetry pathway in K-to-DVB conversion and suggest that the wild type orientation of K division is not a requirement for DVB formation from K.p.

### Both selection of one K daughter and the K-to-DVB conversion require WNT activity

While we found that the orientation of K division *per se* does not impact on DVB formation, this did not preclude a potential role for the Wnt pathway in discriminating the K daughters. Indeed, the Wnt/β-catenin asymmetry pathway is also known to control the polarity of several cell divisions during *C. elegans* development through the WNT ligands (Herman, 2002; Herman et al., 1995). We therefore examined the phenotypes of the posterior WNT ligands *lin-44/WNT* single and *lin-44/WNT; egl-20/WNT* double mutants (Fig. 3B-C) in greater detail. *lin-44* is expressed posteriorly to K in tail hypodermal cells, while *egl-20* is expressed in some rectal cells, including K, and in other cells found in the rectal area (Harterink et al., 2011). The DVB neuron is present in most of the single *lin-44* and the double *lin-44; egl-20* mutant animals (Fig. 3B-C), contrary to what observed in *lin-17* mutant. However, we found that DVB appears to originate from the K.a daughter in 10.8% of *lin-44* mutant worms and in up to 57.9% in animals lacking also *egl-20* (Fig. 3C), a reversed polarity of cell division phenotype as observed in T and the male B, F and U cell divisions (Herman et al., 1995). Moreover, consistently with a role of WNTs acting as positional cues and determining the polarity of the division outcome, we observed that LIN-17/FZD receptor transiently localizes at the posterior cortex around 2h before cell division (not shown). By contrast, reversed polarity of cell division is never observed in in Wnt pathway mutants downstream of WNTs: in both *pop-1/TCF* and *lin-17/FZD* mutants K division always occurs and, in those few instances where a DVB is made, it is from the K.p cell (not shown). We conclude that the Wnt/β-catenin asymmetry pathway regulates both the polarity of K division and K.p conversion into a DVB neuron.

To dissect how this latter process occurs, we characterized K.p identity in *lin-17* mutant as previously done in the wild type. First, K.p nucleus appears as big as K.a nucleus at all times in 77% of *lin-17(n671)* mutants, reminiscent of a hypodermal identity (Fig. 2E). Moreover, all the examined rectal and epithelial markers remain expressed in K.p, even after it should have already become DVB: this is the case for continued *let-413* and *lin-26* expression (Fig. S4B-C, Table S1), and even for *ajm-1*, which is found in K.p although it does not inherit this K apical junction protein (Fig. S4A, S5, Table S1). Similarly, all rectal markers tested, such as *egl-5/HOX* are retained in K.p in *lin-17* mutant at all stages (Fig. S4D, Table S1). On the contrary pan-neuronal (*rgef-1/RASGRP3, unc-119/UNC119* and *unc-33/DPYS*) and GABAergic markers are never expressed (Fig. S4E-I, Table S1). These findings suggest that the K posterior daughter K.p retains a rectal identity in the absence of *lin-17*.

In sum, K-to-DVB involves several steps during which the Wnt/ β-catenin asymmetry pathway plays at least two distinct roles: i) *lin-44/WNT* is necessary to assign which of the two K daughters will subsequently change into a DVB neuron, probably acting as a positional cue coming posteriorly to K; and ii) Wnt signal transduction allows the K.p cell to ultimately convert into a DVB neuron. Furthermore, K division *per se* appears necessary, but not sufficient, for the ability of the K cell to ultimately produce a DVB neuron.

### Key Y-to-PDA plasticity factors are also required for K-to-DVB conversion

We had previously demonstrated that *sem-4*/*SALL, egl-27*/*MTA, ceh-6*/*POU* and *sox*-2/*SOX* are necessary for the erasure of Y rectal identity during Y-to-PDA Td and that *egl-5/HOX* is a likely downstream effector during Y dedifferentiation (Kagias et al., 2012). Since our data suggest the requirement, reminiscent of Y-to-PDA, of mechanisms additional to K division, we investigated whether Y “plasticity cassette” genes could be required for K-to-DVB conversion as well. Supporting this possibility, traditional and our CRISPR-KI reporters show that these factors are also expressed in K (Table S1, Ferreira et al., 1999; Bürglin and Ruvkun, 2001; Jarriault et al., 2008; Vidal et al., 2015).

We found that *sem-4(n1971)* and *egl-5(n945)*, which are viable null mutants (Chisholm, 1991; Basson and Horvitz, 1996), display the strongest DVB defect with respectively 92% and 85% “NO DVB” worms (Fig. 4A), a penetrance similar to the PDA defect observed in these mutants (Jarriault et al., 2008). As absence of *ceh-6* and *sox-2* is lethal before the conversion occurs, we assessed their involvement using a rectal-specific mutant for *ceh-*6*(gk665)* (Ahier et al., 2020) and rectal-specific mild RNAi or anti-GFP nanobody strategy for *sox-2* (Wang et al., 2017). *ceh-6(gk665)* and *sox-2* knockdown showed significant defects in PDA and DVB formation (Fig. 4A), with a low penetrance most probably due to experimental limitations (Vidal et al., 2015). To rule out involvement of other family members in light of the low penetrance in *ceh-6* and *sox-2* mutants, we also tested all their paralogs (Fig. S6), but none displayed any significant “NO DVB” defect, confirming that *ceh-6* and *sox-2* are the core components required for cell fate conversion. Consistently, Vidal and colleagues had found that the *sox-2* paralog *sox-3* is not required for DVB formation, even though co-expressed with *sox-2* in rectal cells (Vidal et al., 2015). Finally, we observed that the absence of neither *egl-27*, nor of the *egl-27* paralog *lin-40*, have a strong impact on DVB formation as opposed to Y-to-PDA Td (Fig. 4A, S6).

**Fig. 4.**
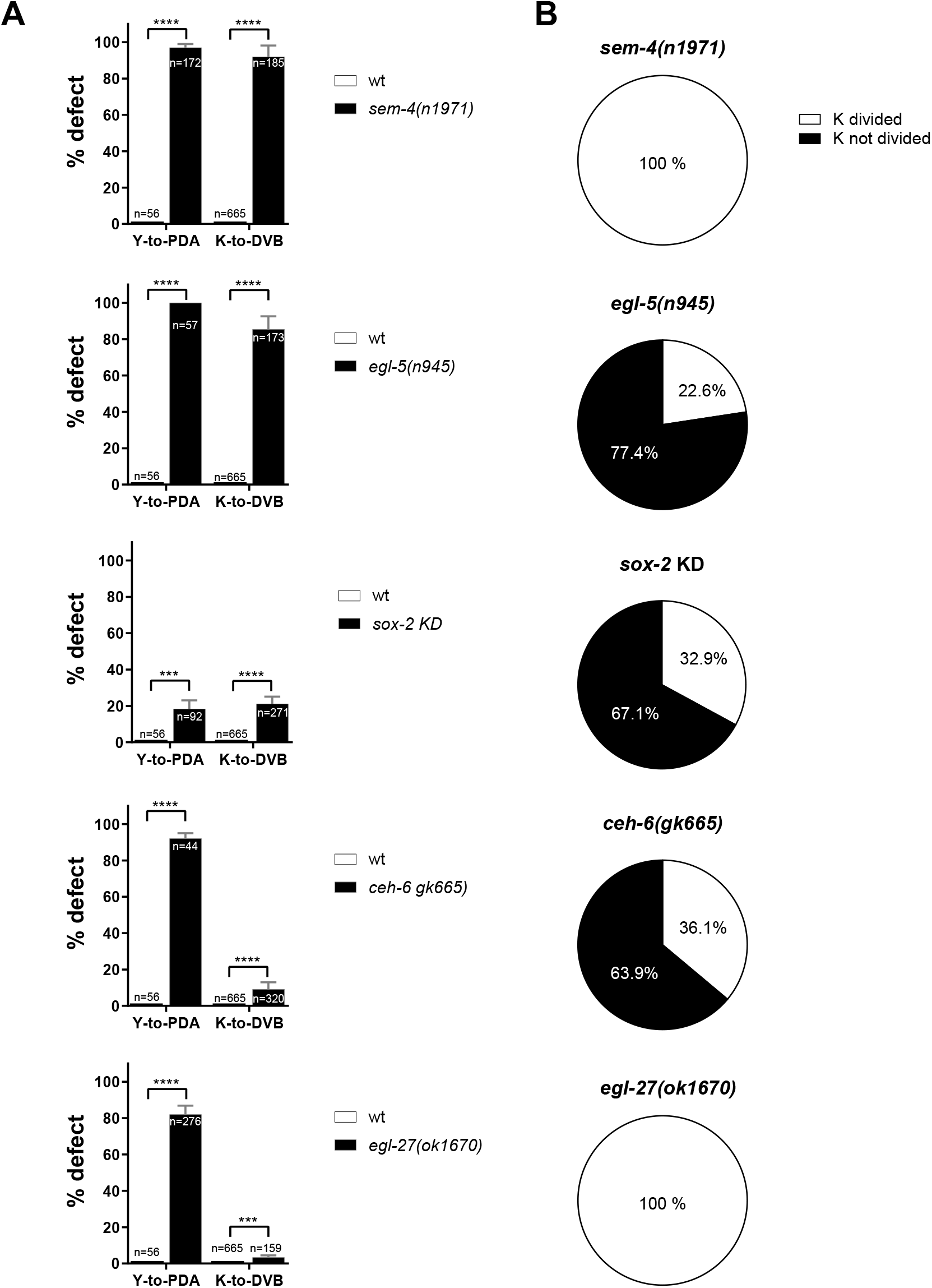
Some key Y-to-PDA plasticity factors are also required for K-to-DVB. **(A)** Quantification of “absence of PDA” (observed with *exp-1p::gfp)* and “absence of DVB” (observed with *unc-47p::gfp*) defects in wild type L4 *vs* mutants or deficient background for Y-to-PDA key plasticity factors. **(B)** Quantification of K division in deficient backgrounds for Y-to-PDA key plasticity factors. 276, 87, 221, 127 and 160 animals were scored in *sem-4, egl-5, sox-2, ceh-6* and *egl-27* mutants, resp.

Altogether these results demonstrate that most of the genes required for Y-to-PDA, namely *sem-4, egl-5, sox-2* and *ceh-6*, in addition to being expressed in K, are also required for K-to-DVB conversion.

### The “plasticity cassette” enables K division and erasure of K.p rectal identity

Since cell division is necessary for DVB formation, we analyzed the K cell division in the *egl-5, sox-2, ceh-6* and *sem-4*-deficient backgrounds. We could not assess the orientation and asymmetry of K division in *egl-5(n945), ceh-6* or *sox-2* mutant backgrounds due to technical limitations (see Methods). However, we could observe that the absence of DVB in *egl-5* null, *ceh-6* rectal-specific mutant and *sox-2* knockdown using the anti-GFP nanobody strategy is due to K not dividing in three fourth (for *egl-*5) and two thirds (for *ceh-6* and *sox-*2) of the cases (Fig. 4B). When K divided, we saw that in worms that failed to develop DVB, K.p retained the rectal identity as visualized by sustained *col-34* expression in L4 larvae (not shown). Thus, these genes appear to have at least two roles during K-to-DVB: i) allow the occurrence of K division; and ii) allow the K.p daughter to undergo an identity change, very reminiscent of their role during Y-to-PDA Td. Conversely, K division occurs in 100% of the animals in *sem-4(n1971)* null mutant (Fig. 4B) and its orientation appears similar to the wild type orientation (Fig. 2D). Since K division looks normal and K.p is always formed but does not give rise to DVB in more than 90% of the animals, we analyzed K.p identity in *sem-4* mutant. We found that K.p exhibits a large nucleus (Fig. 2E) reminiscent of an epithelial identity, in agreement with previous DIC observations (Basson and Horvitz, 1996), a phenotype that persists to the adult stage. We further found that K.p retains the epithelial and rectal markers *col-34, lin-26* and *let-413* in L3 and L4 *sem-4(n1971)* mutant larvae, when K.p has already become DVB in wild type animals (Fig. S4B-C and Table S1); most importantly we also observed *ajm-1* expression, suggesting that this gene is never silenced or is re-expressed in K.p in *sem-4* (Fig. S4A, S5 and Table S1). In addition, we found persistent expression of all rectal markers analyzed in those mutant K.p, while they disappear from the wild type K.p (e.g. *egl-5* and *col-34*, Fig. S4D and Table S1). On the contrary, DVB neuronal markers, including pan-neuronal (*rgef-1, unc-119, unc-33*) and GABAergic markers (*unc-47* and *unc-25*), are not expressed in K.p in *sem-4* mutant (Fig. S4E-I and Table S1). We conclude that in absence of *sem-4*, like in *lin-17* mutant, while K division appears normal, K.p cannot eliminate its rectal identity which is retained indefinitely.

Collectively, our findings suggest several roles for the plasticity TFs’ genes in K-to-DVB conversion: *egl-5, ceh-6* and *sox-2* appear to act both before and after K division, while *sem-4* activity is required to initiate K.p reprogramming through the erasure of its rectal identity. In addition, our results demonstrate that while K division is required for K-to-DVB, it is not sufficient, in absence of key TFs, to generate a new identity in one of the two K daughter cells, and events involving *sem-4* and other Y-to-PDA “plasticity cassette” factors are also required.

### Plasticity factors and the Wnt signaling pathway act in parallel to erase K.p rectal identity

The requirement of both Y-to-PDA factors and the Wnt signaling pathway for K.p reprogramming prompted us to investigate whether those players act in parallel or in the same pathway. We focused on *sem-4* mutant and its relationship with the Wnt signaling pathway, as *sem-4* and *lin-17* mutant worms show very similar phenotypes. We hypothesized that the Wnt pathway could control *sem-4* expression in K.p. Indeed, it has been demonstrated that TCF/LEF1 can bind to the *SALL4* promoter in human cell lines *in vitro* (Böhm et al., 2006). However, we observed that the expression of a *sem-4* endogenous KI-reporter is not affected in K.p nor in the other rectal cells of *lin-17(n671)* mutant (Fig. S7A). Thus, *sem-4* is not downstream of the Wnt signaling pathway in K.p. In support of this conclusion, we found that a *sem-4* translational construct able to rescue *sem-4* phenotype was not capable to rescue DVB defect of *lin-17* mutant (data not shown), excluding that *sem-4* activity would act downstream of *lin-17*. We tested the reciprocal relationship by building a *lin-17* transcriptional reporter and observed the same *lin-17* faint expression in K.p and in K.a in *sem-4(n1971*) or the wild type (not shown). However, while the expression disappears as it becomes DVB in the wild type, it is retained in the mutant K.p (Fig. S7B). This is consistent with maintenance of the rectal identity of the K.p cell in *sem-4* mutant, as *lin-17* is expressed in the rectal cells. In support of this hypothesis, we found that *lin-17* expression in *lin-17* mutant also persists in K.p, which appears to retain a rectal identity as in *sem-4* mutant (Fig. S7B).

We next examined the genetic relationship between the Wnt signaling pathway and the “plasticity cassette” factors acting on K.p identity using double mutants and analyzing their impact on DVB formation. We used the *wrm-1*/β-catenin *(n1982ts)* mutant for the Wnt pathway since we could not use *lin-17/FZD* mutant due to its high penetrance of DVB absence. For the same reason, we chose to use the *sem-4* hypomorphic allele *n1378* that shows a low “NO DVB” defect (Fig. S7C) rather than null mutant *sem-4(n1971)*. We found that the double mutant *sem-4(n1378)*; *wrm-1(n1982ts)* led to a synergistic “NO DVB” defect compared to the single mutants at 25°C (Fig. S7C). Confirming this genetic relationship between the Wnt pathway and plasticity cassette factors, we furthermore observed the same synergy in double mutants with downregulated *sox-2* and *wrm-1* activities (Fig. S7D).

Altogether our results suggest that the plasticity factors and the Wnt signaling pathway act through two different parallel genetic pathways to control the loss of K.p rectal identity. Y-to-PDA and K-to-DVB require the same set of TFs, independently of the activity of the Wnt pathway and cell division occurrence, and thus appear to share common cellular and molecular mechanisms to remove the initial identity, providing further evidence that K-to-DVB is a bona fide Td event (Fig. 6A, B).

### Antiparallel activities of *sox-2/ceh-6* and the Wnt signaling may control the timing of re-differentiation

After having identified the factors required for the loss of K.p rectal identity, we wanted to dissect the mechanisms regulating re-differentiation into DVB. To this end, we focused on *lim-6* gene, encoding a LIM homeobox TF and the sole DVB terminal selector identified to date: in *lim-6* mutant, DVB terminal differentiation is not complete (Hobert et al., 1999). First, we determined the expression timing of *lim-6*. Using both a translational rescuing construct and a CRISPR-KI allele that we engineered, we observed that *lim-6* starts to be expressed in K.p in around 25% L1 wild type larvae 1 to 2 hours after K division (Fig. 5A), overlapping with the decreasing expression of *lin-26* and *let-413* epithelial markers (Fig. S2B). A fragment corresponding to the large intron 4 in the *lim-6* locus was found to be sufficient for the expression of a reporter gene in DVB (Hobert et al., 1999). We built a similar transcriptional reporter bearing the intron 4 fused to *gfp* and we found that it follows the same transcriptional dynamics in K.p as the KI reporter (not shown). Thus, *lim-6* expression appears to be an early indicator of K.p conversion and DVB future identity (Fig. 5A, Fig. S2B). We next examined its expression in *sem-4* and *lin-17* mutants using both the intron 4 transcriptional reporter and the CRISPR-KI: we observed an absence of *lim-6* expression in K.p. (Fig. 5B, S9B and data not shown), confirming that K.p remains a rectal cell in those mutants.

**Fig. 5.**
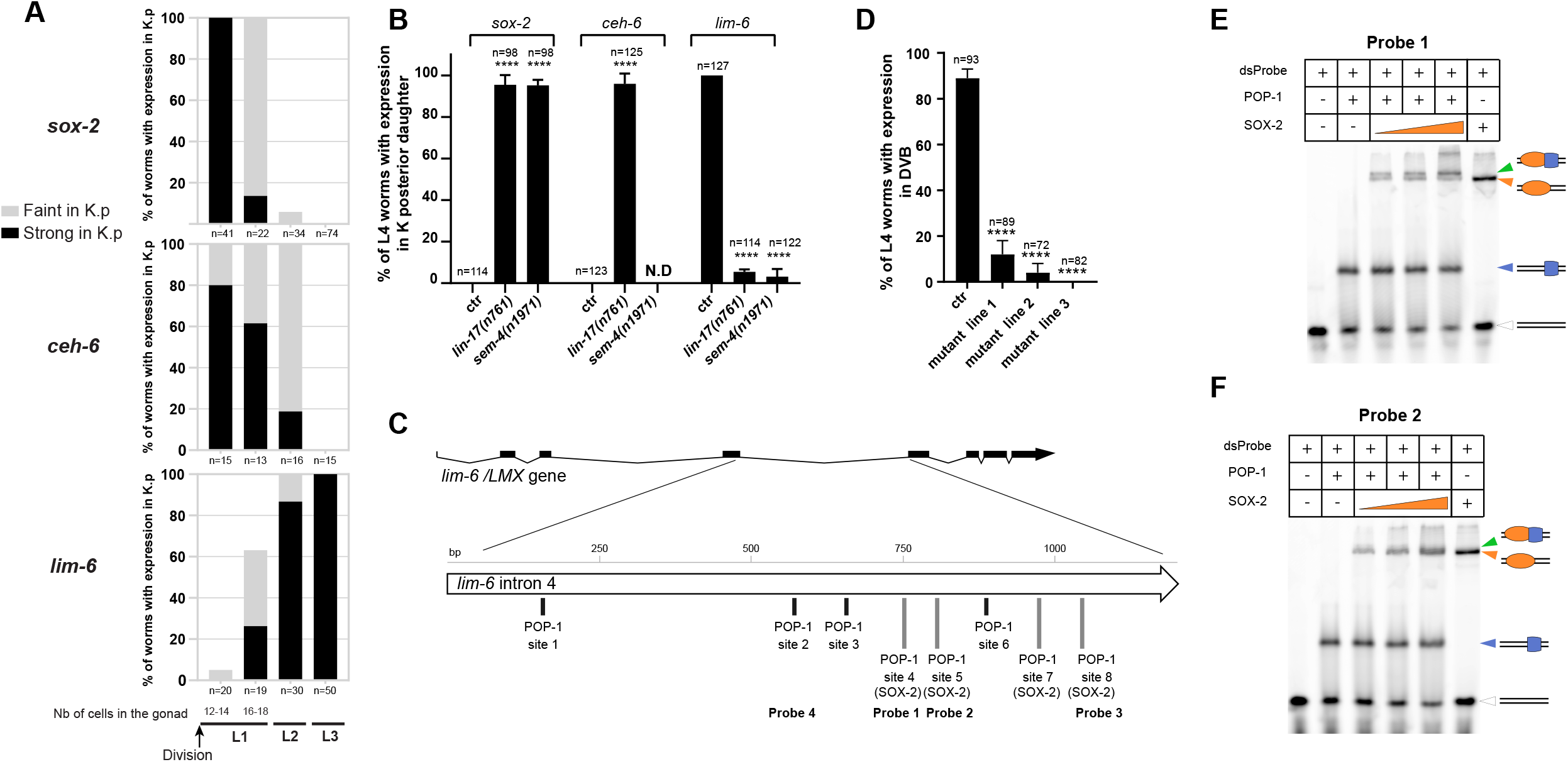
lim-6 expression regulation by SOX-2 and POP-1/TCF. **(A)** Time course expression of *sox-2, ceh-6* and *lim-6* CRISPR-KI reporters in K.p right after K division in L1, in early L2 and in L3 (DVB). For L1s, worms were tightly synchronized based on the number of cells in the gonad. Black bars represent strong signal in K.p, grey bars represent faint signal in K.p. **(B)** *sox-2, ceh-6* and *lim-6* CRISPR-KI reporters expression in L4 in wild type (DVB), *lin-17(n761)* and *sem-4(n1961)* mutant backgrounds (persistent K.p). *ceh-6* CRISPR-KI expression could not be addressed in the *sem-4* mutant due to genetic linkage of both genes. **(C)** Genomic organization of *lim-6/LMX* and POP-1/TCF and SOX-2 binding sites in the 4^th^ intron. The probes used in this study are represented. **(D)** Expression in DVB of the *lim-6* intron 4 transcriptional reporter depends on TCF sites. One transgenic line bearing the wild type version of the intron 4 and three independent transgenic lines bearing the mutated intron 4 (TCF sites 2 to 8) were analyzed in parallel in L4 larvae. (B, D), Bars represent mean +/-standard deviation. n, total animals scored. (**E-F**) Gel shift assays using the purified full-length SOX-2 and HMG-POP-1 with CY-5-labeled double stranded DNA probes 1 or 2. Increasing concentrations of SOX-2 (125nM-250nM-500nM) were added to the dsProbe and HMG-POP-1. Binding of HMG-POP-1 was observed on both probes in the absence of SOX-2 (blue arrowhead). Increasing quantities of SOX-2 translate into increased binding to the probes (dark orange arrowhead) as well as an upper shift (green arrowhead) most probably corresponding to a HMG-POP-1-SOX-2-Probe complex. Double line, free probe; double lines with blue square, orange oval or both represent the HMG-POP-1-Probe, the SOX-2-Probe and a SOX-2-HMG-POP-1-Probe complex, resp.

**Fig. 6.**
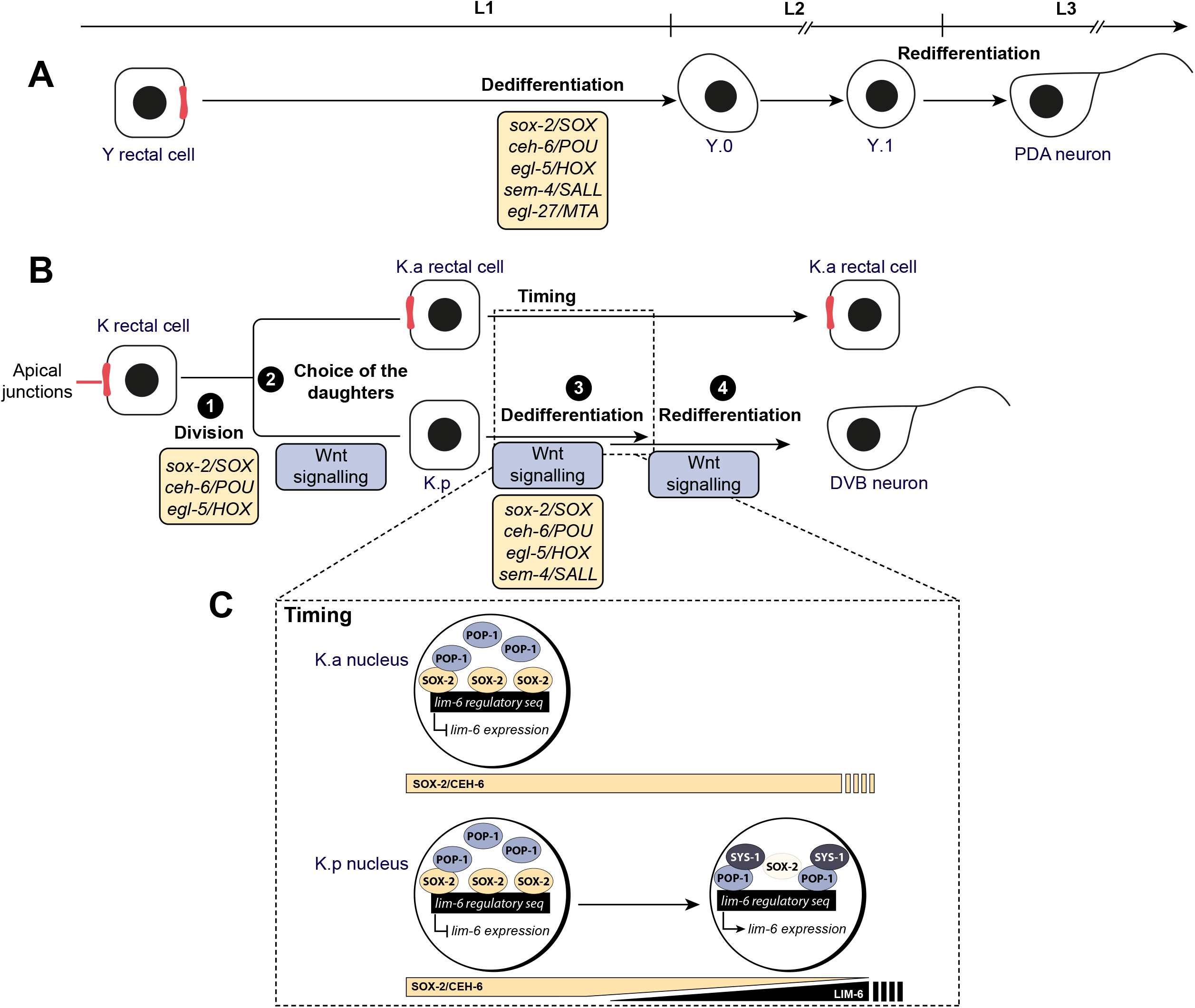
Model for K-to-DVB Td and parallel with Y-to-PDA. **(A)** Y-to-PDA Td, which is initiated at the end of the L1 stage and finalized in the L3 stage, goes through two intermediate states: Y.0, which appears to have lost characteristics of the initial identity but not have gained those of the final identity; and Y.1, which appears to be an early neuronal cell. The box describes the cell plasticity cassette factors required for the initiation of Y-to-PDA Td, that is the dedifferentiation step. Top: developmental timeline for (A) and (B). **(B)** K-to-DVB Td, which is initiated towards the end of the L1 stage and finalized in the L2 stage, involves a cell division and an intermediate state that possibly represents a mixed identity between the initial and the final identity. The four important features of this Td, as well as the factors involved, are highlighted: division, choice of which daughter will adopt a different identity, loss of the initial identity and adoption of a subtype-specific final identity. **(C)** Inset representing how the dynamics of SOX-2 and POP-1 levels and their competition in modulating *lim-6* expression can provide a timer for the re-differentiation step.

The earlier expression of *lim-6* in K.p compared to the other neuronal markers led us to hypothesize that it could be directly downstream of the Wnt/β-catenin asymmetry pathway, through POP-1 binding to *lim-6* regulatory regions. Interestingly, studies in mouse demonstrated that *Lmx1*, the *lim-6* ortholog, is regulated by the Wnt signaling in proliferative dopaminergic neuron progenitors (Joksimovic et al., 2009, 2012; Chilov et al., 2010). We looked for putative POP-1/TCF binding sites in the intron 4 of *lim-6* using several platforms and manual inspection (see Methods) and found 8 sites conserved across several nematode species (Fig. 5C and S8A). When *gfp* reporter constructs lacking the regions encompassing sites #3 to #6 or #3 to #8 were injected in the worm, expression was absent in DVB (as well as in the PVT, AVL and RIS neurons, data not shown). To test whether those sites are specifically required for expression during the K-to-DVB Td, we eliminated them by targeted mutations (Arata et al., 2006), focusing on the well conserved sites #2 to #8. Transgenic animals bearing the mutated constructs showed almost no expression in DVB (Fig. 5D), suggesting that *lim-6* is indeed directly, positively regulated by the Wnt signaling during K.p-to-DVB conversion.

Interestingly, POP-1/TCF binding sites may overlap with SOX2 binding sites, since both TCF and SOX2 are HMG domain proteins (Lin et al., 1995; Pevny and Lovell-Badge, 1997). Indeed, we found that 4 of the POP-1/TCF binding sites in the intron 4 of *lim-6* overlap with SOX2 binding sites (Fig. 5C, S8A and Methods). This observation prompted us to characterize the relationship between SOX-2 and POP-1 and their genes in K.p. We observed that the rectal markers *sox-2* and *ceh-6* (Kagias et al., 2012) display opposite expression dynamics with respect to *lim-6* in K.p (Fig. 5A), resulting in a complete absence of expression in DVB (Table S1). The early onset of *lim-6* expression in K.p and its anti-correlation to *sox-2* and *ceh-6* expression led us to hypothesize that a competition between SOX-2(/CEH-6) and POP-1 could result in the *lim-6* expression timing that we observe. To explore a possible antagonistic function of POP-1 and SOX-2 on *lim-6* expression at the molecular level, we examined if both POP-1 and SOX-2 could bind to *lim-6* intron 4 sequences. Electrophoresis mobility shift assay (EMSA) experiments using DNA probes corresponding to TCF/SOX2 binding *sites #2, 4, 5* and *8* in *lim-6* intron 4 (Fig. S8) showed that both SOX-2 and HMG-POP-1 can bind to these sequences alone (Fig. S10 and not shown) and together (Fig. 5 E-F, S11). Thus, it is possible that SOX-2 and POP-1 display antagonistic activities on *lim-6* activation depending on their expression levels and nuclear localization over time. The dynamic expression of *sox-2* (Fig. S9A) and the nuclear localization of POP-1/β-catenin might further reflect on the precise developmental timing of the expression of *lim-6* gene (Fig. 6C). Interestingly, *sem-4* and *lin-17* appear to be required for *sox-2* and *ceh-6* downregulation in K.p (Fig. 5B), connecting the erasure of the rectal identity by *sem-4* and the Wnt signaling pathway and the re-differentiation into the DVB neuron which probably requires *sox-2* and *ceh-6* downregulation.

In sum, our results suggest that the Wnt signaling pathway and Y-to-PDA plasticity cassette factors contribute to both loss of the rectal identity and re-differentiation into DVB, directly and indirectly: *sem-4* and *lin-17* activities lead to the loss of rectal identity and thus downregulation of *sox-2* and *ceh-6* rectal genes in K.p, where the latter might have a repressive function on *lim-6* expression. In parallel, the activation of the Wnt signaling pathway likely triggers the activation of *lim-6* through direct binding of POP-1, possibly in complex with the SYS-1 β-catenin, to *lim-6* regulatory regions (Fig. 6C).

## Discussion

This study aimed at characterizing the requirements for a natural reprogramming event occurring through a cell division during *C. elegans* larval development: the formation of the DVB GABAergic neuron from the K rectal cell. Indeed, we have found that the K rectal cell, that forms an integrated structural part of a vital organ, the rectum, is a fully differentiated and specialized cell, with all characteristics of the end-of-lineage rectal identity. Yet, this differentiated cell divides once. Few other terminally differentiated cells have been described as able to divide, like hepatocytes which form similar hepatocyte daughters (Miyaoka and Miyajima, 2013). The rectal K cell, by contrast, gives rise to 2 daughters, one which subsequently adopts a different, differentiated, identity. This very interesting process is reminiscent of a stem cell behavior associated with cellular plasticity, with 2 major differences: only 1 round of division is ever observed, and the mother cell that divides is unambiguously differentiated, in fact so similar at the transcriptomic level to the other five non-dividing differentiated rectal cells that all rectal cells cluster as one (Packer et al., 2019). The ability of a differentiated cell to give rise to another differentiated cell has been termed transdifferentiation (Eguchi and Kodama, 1993; Lambert et al., 2021) and K-to-DVB meets the three criteria defining a Td. Since other natural direct reprogramming events can occur in absence of cell division across phyla (Jarriault et al., 2008; Richard et al., 2011; Molina-Garcıa et al.; Karow et al., 2018), this raises the question of the impact of the cell division and of possible differences in underlying mechanisms. In probing these mechanisms, we have shown that several factors, including cell division, conserved reprogramming TFs and the Wnt signaling pathway, are required for this transdifferentiation event at different steps, highlighting common players and alternative mechanisms with Td events involving no cell division.

### Cell division is required for K-to-DVB and may contribute to reprogramming through different mechanisms

Using mutants defective for K division we have demonstrated that cell division is a step required for DVB formation. How might it contribute? Since K division is oriented, one mechanism might involve the possible asymmetric partitioning of cellular components. However, our data do not point to such mechanisms: plasticity cassette factors SOX-2, CEH-6, SEM-4 or EGL-5, the epithelial determinant LIN-26 or any of the rectal markers are not asymmetrically partitioned (we could not examine this for POP-1/TCF and the β-catenins BAR-1 and SYS-1 since their expression levels are too low in the K cell and its daughters). Of note, some epithelial features remain only in the anterior daughter K.a, like the asymmetrically distributed apical membranes and possibly some associated proteins. Nevertheless, while absence of the E-cadherin facilitates the differentiation of pluripotent stem cells into neural cells (Malaguti et al., 2013), the anterior daughter of K can still turn into a DVB-like cell in Wnt mutants (this study), suggesting that this is not the case in this process. Future studies aimed at exploring the importance of apical membranes for K cell daughters’ identity will be warranted to further rule out this hypothesis.

Another possible key contribution of cell division could be the occurrence of DNA replication (Nashun et al., 2015). This might facilitate the erasure of pre-existent chromatin marks and/or provide accessibility to portions of the genome that would otherwise be in a closed conformation in the K rectal cell. In addition, studies of induced direct reprogramming suggest that chromatin regulators can act as barriers to reprogramming (Hajduskova et al., 2019; Kolundzic et al., 2018; Rothman and Jarriault, 2019). However, our analysis of the *lin-5* mutant in which cell division fails at the cytokinesis step suggests that DNA replication alone is not sufficient to trigger a dramatic transcriptional change and the erasure of K rectal identity. This does not rule out a role for DNA replication in facilitating the Td, for instance by dispensing with some of the factors required for reprogramming events in which DNA replication does not occur. In support to this, we have found that *egl-27/MTA*, an ortholog of a chromatin remodeling complex component (Kumar and Wang, 2016), is not key for K-to-DVB conversion while it is very significantly required for Y-to-PDA Td which occurs in absence of cell division (Kagias et al., 2012).

On the more speculative side, the generation of an additional cell to make a DVB motor neuron might be under positive selection pressure since the K cell is a non-redundant structural part of the rectal tube, a vital organ (Sulston et al., 1983; this study). In support to this hypothesis, the Td of the rectal Y cell (Jarriault et al., 2008), or of the phasmid socket PHso1 (Molina-Garcıa et al., 2020), which occurs in absence of cell division, also occurs while other cells, P12.pa and PHso2 respectively, will replace functionally the transdifferentiating cell. By contrast, no other cell replaces the AMso amphid socket, which conversion involves a cell division - although its importance for the cell conversion is not known (Sammut et al., 2015). It is intriguing to note that known examples of “no division” Td events involve a retraction process similar to an Epithelial-to-Mesenchyme-Transition (EMT, Yang et al., 2020). EMT-like process and cell division might serve the same purpose each time, facilitating the removal of the initial specialized epithelial identity.

### The Wnt/β-catenin asymmetry pathway mediates several aspects of K-to-DVB Td

Our data support the involvement of the *C. elegans* Wnt/β-catenin asymmetry pathway during K-to-DVB. The Wnt/β-catenin asymmetry pathway influences cell identity specification in several cellular contexts and at different developmental stage in *C. elegans*, ranging from the 2-cell stage, to late embryonic development and to larval development (Mizumoto and Sawa, 2007; Bertrand and Hobert, 2009; <u>Wormbook</u>). Our study shows for the first time that this pathway can also contribute to Td of a fully differentiated cell in multiple ways: the choice of which daughter will convert to a different identity; the erasure of the rectal identity; and the definition of DVB subtype-specific identity (Fig. 6B).

When a differentiated cell divides, what mechanisms ensure that one of the daughter cells only subsequently adopts a different identity? Two non-exclusive mechanisms can generate an asymmetry in daughter cells’ fate : asymmetric partitioning of cell components (molecules, organelles, etc), or providing similar daughters with a different environment leading them to adopt different fates (Betschinger and Knoblich, 2004). Here we show that an exterior signal, possibly combining with physical constraints (see below), ensures that one daughter is formed at a stereotyped posterior-basal position and is able to lose its mother identity. The phenotypes of two different WNT ligand mutants synthetized by cells in the tail region, suggest that the ligand important for K-to-DVB is LIN-44, and that, in absence of LIN-44, other anteriorly expressed WNT ligands can induce K-to-DVB, as observed in seam cells (Yamamoto et al., 2011; Jackson and Eisenmann, 2012). This would explain both the non-fully penetrant “NO DVB” defect in the *lin-44* mutant, and the reversed polarity of cell division observed in this background. Thus, we propose that the Wnt signaling pathway has an instructive, rather than a permissive role on K-to-DVB Td, with WNT ligands acting as positional cues in this process, instructing which K daughter will cease to be a rectal cell. An analogous situation has been recently demonstrated in the SMDD/AIY asymmetric cell division during embryonic development (Kaur et al., 2020).

The stereotyped division orientation could contribute in selecting K.p for a cell identity conversion. For instance, it could ensure that the posterior daughter sees more, or more rapidly, of the posterior WNT ligand than K.a, ensuring no disruption of the rectum function. Activation of the Frizzled receptor by a WNT ligand in dividing cells was shown to impact on the orientation of the mitotic spindle at least in the early blastomeres (Schlesinger et al., 1999). Consistently, defects in the orientation of K division are observed in *lin-17* mutant as well as transient localization of LIN-17/FZD at the posterior cytoplasmic membrane in K just before its division (not shown). However, while the impact of *lin-17* null mutation on DVB formation is almost fully penetrant, the percentage of K division orientation defect is much lower. One possibility is that not only the Wnt signaling pathway, but also physical constraints may regulate the orientation of K division. The K cell is integrated in the rectal tissue, forming apical junctions (the *C. elegans* equivalent of both adherens and tight junctions; Armenti and Nance, 2012) with the neighboring rectal cells. Such tissue structure appears maintained during K division. We propose that those features of K and the rectal tissue might impact on the orientation of the division together with the Wnt pathway and ensure its robustness, similarly to the role of cell shape and the Wnt pathway described by Wildwater et al., 2011. This might explain why the mutants that we have tested (*gpr-1, goa-1* and *gpa-16* mutants), have not shown any significant impact on K division orientation, by contrast with their impact on spindle orientation in early fast-dividing blastomeres that lack adherens junctions and organ building function (Nance and Priess, 2002).

Irrespective of the division orientation, in absence of the Wnt signaling the K.p cell remains a rectal cell that does not exhibit any neuronal features or expression of neuronal markers, suggesting that one important role of the Wnt signaling is to allow the K.p cell to shed its epithelial identity, reminiscent of Y dedifferentiation step. A role for this pathway has recently been proposed in promoting other Td events: in hepatocyte-to-cholangiocyte Td (Kosar et al., 2020), and in the conversion of biliary cells (BEC) into hepatocytes in mouse organoid models and injured liver where hepatocyte proliferation is compromised (Valle-Encinas et al., 2020). It would be interesting to assess if the WNT signal results in promoting the erasure of the initial identity in these instances as well. Furthermore, BEC-to-hepatocyte conversion in chronic injury human liver models had been shown to involve intermediate states exhibiting BEC and hepatocyte markers (Deng et al., 2018), suggesting a transition through a mixed identity, as our markers analysis suggests for the K.p-to-DVB transition. Thus, although the exact function of the Wnt pathway remains to be elucidated in these contexts, it might be that the mechanisms by which WNT impacts on direct cell identity conversions are conserved.

Interestingly, the Wnt pathway acts not only to promote dedifferentiation, but also re-differentiation, through the acquisition of DVB-specific identity. We found that the Wnt pathway is required for expression of the LIM-6 TF, determining which neuron subtype will be formed. LIM-6 is considered as one of DVB terminal selectors, that is, a TF responsible for the induction and maintenance of a battery of terminal differentiation genes (Hobert, 2016). This is evidenced in *lim-6* mutant, where the DVB neuron is formed but lacks part of its terminal features (Hobert et al., 1999). We cannot easily assess the dynamics of K.p dedifferentiation and re-differentiation into a DVB neuron and whether WNT distinct functions are sequential. In support of sequential outputs however, it is interesting to note that WNT signal has been shown to have opposite effect on ES cells depending on the cell state (Merrill, 2012). Future studies, for instance looking at the evolution over time of the transcriptome of the K.p cell, may address this fascinating question.

### Conserved reprogramming transcription factors are required for K-to-DVB

We have shown that the Y-to-PDA factors SEM-4/SALL, EGL-5/HOX, SOX-2 and CEH-6/POU are also required for K-to-DVB to erasing K.p rectal identity after the division, and also for K division to occur (the latter three). It is striking that these factors are involved similarly to their function in Y, further reinforcing the notion that shared mechanisms exist (Fig. 6A, B). Importantly, our findings strongly suggest that the division of K, which we showed to be necessary to the formation of a DVB neuron, is not sufficient in itself to produce a K daughter with a neuronal fate: in *sem-4* mutant, the K cell division occurs mostly like in wild type; however, the K.p cell fails to activate the expression of neuronal genes, and remains stably there, displaying a large, epithelial-like nucleus, as previously described in (Basson and Horvitz, 1996), and expressing epithelial and rectal genes as the other rectal cells. Reinforcing this conclusion, in *sox-2* and *ceh-6* mutants where the K cell divided, the perdurance of *col-34* expression in K.p, associated with the absence of DVB, similarly suggests that K.p also remains rectal. We note that our data are in agreement with the previously postulated role for *sox-2* for the terminal neural differentiation of post-embryonic differentiated epithelial cells, aka transdifferentiation (Vidal et al., 2015), with a difference: Vidal and colleagues conclude that the K.p cell found in *sox-2* mutants does not acquire rectal identity like its sister K.a based on lack of *ceh-6* expression (used as rectal cell marker), contrary to what *col-34* expression as observed here suggests. However, the observed lack of *ceh-6* expression in K.p in *sox-2* mutants could be due to the continued activity of the Wnt signaling pathway as we found it to be required for *ceh-6* downregulation, rather than reflecting a loss of rectal identity. In addition, the K cell appears able to enter the cell cycle in absence of *sox-2* (Vidal et al., 2015) although we see a significant proportion of worms where the K cell fails to divide, suggesting that the K cell division is probably initiated in most animals but fails in a significant portion of them, notably leading to a cell with two separate sets of segregated sister chromatids.

Overall, these results have led us to hypothesize that those factors might be part of a conserved “plasticity cassette” in *C. elegans*, allowing reprogramming initiation of differentiated cells. Interestingly, these genes belong to families known to be master regulators of cell fate and their reprogramming capacity is conserved through evolution (Julian et al., 2017; Malik et al., 2018; Tsubooka et al., 2009). We speculate that the conserved reprogramming capacity of the *C. elegans* SOX-2 and CEH-6 in Y and K could be due to pioneer activity as shown for the mammalian counterparts (Soufi et al., 2015). It remains to be seen if these factors are involved in other natural reprogramming events, or if region or tissue-specific variations exist. Interestingly, a recent study reported a low penetrance defect in mosaic animals having lost a *sox-2* rescuing array, although using systemic RNAi the *sem-4* and *sox-2* genes appeared little or not involved, in the AMso-to-MCM or the PHso-to-PHD conversions (Molina-Garcıa et al., 2020). This study together with the present one underscore the importance of the approach used to eliminate *sox-2* activity, since we found that *sox-2* systemic RNAi, even in a sensitized background, did not result in significant K-to-DVB defects by contrast with cell-specific expression of a *sox-2* antisense. In addition, it is conceivable that variations around this set of factors, with the involvement of other family members, such as *sox-3*, for instance, exist between Td events.

It is noteworthy that these plasticity cassette genes are expressed in all rectal cells, including those that never change their identity. While the Wnt pathway has an instructive role on K-to-DVB, our data are consistent with the TFs *sem-4, sox-2, ceh-6* and *egl-5* providing a permissive cellular context for reprogramming in K, as we previously proposed for Y-to-PDA (Kagias et al., 2012). We anticipate that other factors, such as ZTF-11(Lee et al., 2019), are expressed in Y and K and specifically allow reprogramming, and/or that some factors are expressed in the other rectal cells which prevent their plasticity and cell cycle entry in hermaphrodites.

### The Wnt signaling pathway and reprogramming transcription factors cooperate to erase K.p rectal epithelial identity and to control the timing of re-differentiation into DVB

Our characterization of the relationship between the Wnt signaling and the reprogramming TFs, SEM-4/SALL and SOX-2/SOX in particular, suggests that they act in parallel to allow the erasure of K.p rectal features and its conversion into DVB. Such cooperation between the Wnt pathway and members of the plasticity cassette has been described in other contexts and appears conserved: for instance, *sem-4/SALL* and the β-catenin *bar-1* cooperate to render six epidermal precursor cells competent to respond to other developmental signals (Eisenmann and Kim, 2000; Grant et al., 2000), and the Wnt signaling was shown to enhance the pluripotent reprogramming capacities of Sox2, Klf4 and Oct4 (Marson et al., 2008).

We found that one likely consequence of their interplay is the timing of expression of the terminal selector *lim-6*. As outlined above, we found that the Wnt pathway is required for expression of the LIM-6 terminal selector TF. Examination of the DVB enhancer in *lim-6* regulatory region revealed several POP-1/TCF consensus binding sites, which we found to be necessary for *lim-6* expression in DVB and bound by the POP-1 protein. In addition, our observations that *sox-2/SOX* and *ceh-6*/*POU* expression is downregulated in K.p, and this in an anti-correlated fashion to the beginning of the expression of *lim-6*, suggest that the Wnt pathway and at least these two members of the plasticity cassette act antagonistically on *lim-6* expression, and hence DVB neuron-subtype identity acquisition. This hypothesis is further reinforced by our preliminary findings that over-expression of SOX-2 in K.p blocks its conversion into DVB. Since the SOX-2 consensus overlaps with the POP-1/TCF consensus, as both these TFs contain an HMG domain for DNA binding (Lin et al., 1995; Pevny and Lovell-Badge, 1997), this could happen through direct binding to *lim-6* cis-regulatory region and a dynamic interaction with POP-1 at those sites. Indeed, we found that SOX-2 is able to bind to several of these POP-1/TCF sites. Several models can be envisioned: for instance, SOX-2 could compete with low levels of POP-1/TCF for binding and/or preclude *lim-6* expression. However, our gels shift experiments provide little evidence for such model and rather show that both proteins can bind at the same time. Alternatively, through co-occupancy at those sites, SOX-2 could negatively impact on POP-1 activity directly or by sequestering co-factors such as β-catenin. Such model is consistent with studies in vertebrates, where SOX factors have been shown to interact with TCF/β-catenin factors in several ways: at the DNA via interaction with β-catenin and/or TCF, or by impacting β-catenin stability (Akiyama, 2004; Kormish et al., 2010; Mukherjee et al., 2020). The negative interactions between SOX-2 and the Wnt signaling could further be reinforced by CEH-6/POU, as mOct4 has been shown to inhibit Tcf/β-catenin stability and transcriptional activity in mES cells and Xenopus (Cao et al., 2007; Abu-Remaileh et al., 2010). As SOX-2 and CEH-6/POU levels go down, this negative interaction between SOX-2 and the Wnt pathway would be released, and *lim-6* expression activated by POP-1/TCF. Finally, downregulation of *sox-2* and *ceh-6/POU* is lost in *lin-17/FZD* mutant, suggesting negative feedbacks. Thus, we postulate that this dynamic *pas-de-deux* between *sox-2* and the Wnt signaling provides a timer for precise *lim-6* expression in K.p, and therefore re-differentiation timing (Fig. 6C).

Our findings that the Wnt signaling impacts both rectal cell identity loss, i.e. dedifferentiation, and re-differentiation through the promotion of DVB-specific neuronal identity resonate with the study of the role of the Wnt signaling in ES cells, where it has been claimed to promote both the self-renewal, and the differentiation, of ES cells (Davidson et al., 2012; Merrill, 2012). These antinomic effects of the Wnt pathway have been proposed to depend on the cellular state of the ES cells receiving the WNT signal: naïve, or primed for differentiation (Merrill, 2012). A model integrating the pro-pluripotency and pro-differentiation activities of Wnt has been proposed recently, postulating that SOX2 levels influence the cell response to the Wnt signaling (Blassberg et al., 2020): high levels would favor a pro-pluripotency activity of TCF/β-catenin, while low levels would favor caudal epiblast differentiation, through a switch in binding from pluripotency genes to caudal epiblast genes (Blassberg et al., 2020). In line with this model, it will be interesting to test whether the diminishing levels of SOX-2 and CEH-6 in K.p allow the WNT signal to switch from an inhibitor of the rectal transcriptional program to an activator of neuron (DVB)-type identity specific genes. In support of a model positing that dosage of one or more key factors translates into a specific cellular context, *sox-2, as egl-5* and *ceh-6*, is expressed in all rectal cells, including those that do not change identity. However, its expression disappears specifically from Y and K.p during Td, so that it is not expressed in the terminal identities (DVB, PDA, this study; D. Isaia and SJ unpublished). Furthermore, we have observed that increasing the dose of *egl-5, sox-2* and *ceh-6* in the rectal cells before Td, when their concentrations are normally decreasing in wild type animals, led to a block in cellular plasticity and reinforcement of the rectal identity (AA, CG, SJ unpublished). It is thus tempting to speculate that a low dose of these genes allows the switch in transcriptional program possibly through a change in binding factors, when a high dose would maybe participate to the maintenance of the rectal identity: the trigger from stable rectal state to transition into another identity would be through the control of their levels.

### The transition path may differ between different transdifferentiation events

It is striking that K-to-DVB Td occurs in a shorter time-window compared to Y-to-PDA Td (less than 6 hours compared to an entire larval stage). The speed of a fast Td such as K-to-DVB might impact on the nature of the cellular transition. Our analysis of the expression of the terminal selector *lim-6* suggests that the discrete cellular steps observed during Y-to-PDA Td, where the epithelial and rectal genes are turned off before neuronal ones are turned on, may be more blurred in this context, and that co-presence of at least one epithelial and one neuronal marker might be observed for a few hours at the protein level. Further single cell level expression analyses of very large sets of markers and intronic smFISH analyses on the pre-mRNAs of epithelial and rectal genes will allow to precise the nature of the cellular transition states.

Indeed, whether such process proceeds through a discontinuous, step-and-go model, or through a smooth continuous model (Lambert et al., 2021), and how this compares to developmental differentiation trajectories remain open questions. Taken together, our data on K-to-DVB (this study) and Y-to-PDA (Richard et al., 2011) suggest that both continuous and discontinuous modes may exist during Td in *C. elegans*. Such diversity is also seen in reprogramming events outside the worm: Transcriptomic analyses of limb regeneration (Gerber et al., 2018), B-to-macrophage cells (Di Tullio et al., 2011; Francesconi et al., 2018), pericytes-to-neuron (Karow et al., 2018), or fibroblast-to-myotube (Cacchiarelli et al., 2018) reprogramming suggest that these processes occur via a discontinuous, step-and-go model. In addition, transition through a developmental progenitor-like state is often preceded by a transition through a distinct state (Di Tullio et al., 2011; Gerber et al., 2018). By contrast, during fibroblast-to-neuron Treutlein et al., 2016 describe a continuous path with concomitant loss of fibroblast genes and gain of neuronal genes, that nevertheless involves transition through a distinct state expressing a subset of neuroprogenitor genes. Future studies with finer granularity in sampling times within the time window considered, and comparing the proximity of such trajectories to developmental differentiation trajectories will further illuminate the range of paths that lead cells to swap identities in various settings.

## Acknowledgements

We are grateful to J. Godin, O. Hobert, A. Alcolei and M. Barkoulas for their insightful comments on the manuscript. Some strains were provided by the Caenorhabditis Genetics Center (funded by the NIH Office of Research Infrastructure Programs [P40 OD010440]). We thank M. Labouesse for reagents; C. Delance for rectal membrane reporter; A. Daulny and the IGBMC protein purification facility; T. Ye for help with FIMO; Y. Schwab for help with the EM data; and C. LLoret-Fernandez for the original cell design of Fig. 6. This work was supported by the ANR-10-LABX-0030-INRT grant, a French State fund managed by the Agence Nationale de la Recherche under the Investissements d’Avenir frame program ANR-10-IDEX-0002-02, including an IGBMC International PhD Programme fellowship tp CR; as well as by a Ligue Nationale contre le Cancer, an ANR (Agence Nationale de la Recherche) CELLSwitch #ANR-13-BSV2-0005, and an ERC CoG (European Research Council Consolidator Grant) PlastiCell #648960 grants awarded to SJ; MH has been supported by a Ligue contre le Cancer postdoctoral fellowship; CG is a University of Strasbourg assistant professor; and SJ is a CNRS research director.

## Authors contribution

Investigation: CR, MH, CG, AA, SJ - Resources: CR, MH, CG, AA - Validation: CR, MH, CG - Visualization: CR, CG - Conceptualization: CR, MH, CG, SJ - Analysis & Formal Analysis: CR, MH, CG, SJ – Data curation: CG - Writing: CR, CG, SJ - Supervision: SJ - Funding acquisition: SJ - Project administration: SJ

## Methods

### Strains

*C. elegans* strains were maintained on agar plates containing NGM growth media seeded *with E. coli* strain OP50 (Brenner, 1974) at 20°C, except for temperature-sensitive strains *(lin-5(ev571ts), wrm-1(ne1982ts), lin-18(n1051ts), par-1(zu310ts)*) which were grown at 15°C. To score the larvae mutant phenotype in temperature-sensitive strains, embryos, staged via an egg-pulse (spanning over 1h), were shifted at the restrictive temperature (25°C) to avoid early embryonic lethality and scored in L2 or L4. The strains used in this study are summarised in Supplementary Table 2.

## METHOD DETAILS

### Construction of *C. elegans* strains

*C. elegans* transgenic strains were created by DNA microinjection in the gonad of young adults (Mello et al., 1991) of the plasmid of interest together with a co-injection marker and pBSK to a final concentration of DNA of 200-250 ng/µl in water.

All the other strains used in this work (see Key Resource Table) have been obtained by crossing existing strains from our lab, other labs (obtained through CGC or directly) or obtained from SunyBiotech for knock-in reporter strains. The presence of mutant alleles was confirmed by PCR genotyping in case of deletions and by PCR + restriction genotyping in case of point mutations (Morin, 2020). Primers used for genotyping are summarised in Supplementary Table 3.

### Plasmid construction

**pSJ553 – *2nls::gfp***. The 2NLS sequence was amplified by PCR from pSJ207 (Kagias, 2012) with primers oCG390/oCG391 and cloned KpnI/XhoI into pPD95.75.

**pSJ559 – *egl-5p(6*.*5 kb)::nanobodyGFP::zif-1***. *nanobodyGFP::zif-1::U54 3’UTR* was amplified by PCR from pOD1988 plasmid (Wang, 2017) with primers oCR073/oCR074 and cloned into pSJ671 AscI/ApaI sites, containing *egl-5p::*Δ*pes10*.

**pSJ567 – *lin-17p::2nls::gfp***. *lin-17p* (6.5kb) was amplified by PCR from genomic DNA with primers oCR155/oCR156 and cloned into pSJ553 HindIII/PstI sites.

**pSJ721.14 – *let-413::gfp::pest***. The *Mus musculus* ornithine decarboxylase PEST sequence (Corish and Tyler-Smith, 1999) was inserted by Megawhop cloning (Miyazaki 2011) into pML801 plasmid (a gift from Michel Labouesse). The *pest* sequence (120 bp) was obtained through annealing of 2µg of oligonucleotides oCG368 and oCG369 in 10mM Tris pH8, 50mM NaCl and 1mM EDTA. The annealing was performed in 50µl for 5’ at 95°C and then ramping down to about 25°C with a rate of -1.5°C/minute. After the annealing, the *pest* sequence was cloned into pJET1.2/blunt (Thermo Fisher Scientific) and subsequently amplified with primers oCG370/oCG371 and cloned into pML801 by Megawhop cloning.

**pSJ722 – *lin-26rectalp::nls::gfp***. The rectal specific promoter of *lin-26* (Landmann, 2004) was PCR-amplified from genomic DNA with primers oCG381/oCG382 and cloned by Megawhop cloning into pPD97.82.

**pSJ739 – *lim-6int4::gfp***. *lim-6 intron 4* was PCR-amplified from genomic DNA with primers oCG444/oCG445 and cloned by Megawhop cloning into pPD95.75.

**pSJ759 – *lim-6int4(mutated)::gfp***. The sequence of the *lim-6 intron 4* with 7 out of 8 mutated TCF binding sites (Supplementary Figure 8) was ordered (ProteoGenix, France) flanked with BbsI and SalI restriction sites. This allowed it to be cloned into pSJ739, replacing the wild type *lim-6int4* sequence.

**pSJ769 – *T7p::6XHis::hmg-pop-1***. The sequence of the HMG domain of *pop-1* was PCR-amplified from genomic DNA with primers oCG556/oCG557 and cloned by Megawhop cloning into pET30a+.

**pSJ6094 - *T7p::6XHis::sox-2 full::HA***. The *sox-2::HA* sequence was PCR-amplified from peYFP-sox-2-HA (Kagias, 2012) with primers psj6094sox-2 F/R and cloned into pET32a+ SalI/Not1 sites.

**pSJ6293 - *egl-5p(1***,***3kb)::sox-2 full antisens***. *egl-5p(1,3kb)* was PCR amplified with primers pLG7F/pLG7R and cloned into pPD122.53 at the SalI/XbaI sites. The full *sox-2* antisense cDNA was PCR amplified with primers BDT950/952 from a mRNA prep and cloned by Megawhop to replace the GFP present in the original L4053 plasmid.

The coding sequences in the constructs were verified by Sanger sequencing.

All plasmids were transformed into DH10β bacteria (Invitrogen™) for DNA amplification or BL21(DE3) (Stratagene) for protein production.

### RNAi/Silencing experiments

RNAi experiments were performed as previously described (Zuryn, 2014). Basically, we sequence-verified clones from the Ahringer library (Kamath, 2003). RNAi was performed by injecting double stranded RNA (dsRNA) directly into L4 worms, which we found more effective to suppress gene expression in the rectal cells when compared to the feeding method. To this end, the insert of each clone was PCR amplified using T7 primers. I*n vitro* transcription was performed using the PCR products as templates with T7 RNA polymerase using the mMESSAGE mMACHINE™ T7 Transcription Kit (Invitrogen™). Single stranded RNA was allowed to anneal to form dsRNA by gradually lowering the temperature of the sample from 65°C. RNAi sensitised *rrf-3(pk1426) II*; *oxIs12[unc-47::gfp]* adults were microinjected with dsRNA and F1 progeny derived from these adults were scored for the presence of DVB. Note that many of the knockdowns tested resulted in significant lethality and that only escapers were scored, possibly biasing the results towards less defects. For *sox-2* rectal-specific knockdown in particular, we used either the rectal expression (under *egl-5(1*.*3 kb)*promoter) of a *sox-2* antisense sequence, or the rectal expression (under *egl-5*(6 kb) promoter) of the nanobody-GFP system (*nanobodyGFP::zif-1;* Wang et al., 2017) in a *sox-2::gfp* CRISPR KI strain. This latter strategy allowed us to monitor GFP::SOX-2 switch-off in parallel to DVB defects.

### Epifluorescence Microscopy

Worms were immobilized on 2% agarose pads using Tricaine 0.4% and Tetramisole 0.04%. Images were captured on a Leica DM6 B microscope with LAS X software and the HAMAMATSU Digital Camera C11440.

### Scoring criteria

Both localisation, DIC appearance and fluorescent markers, using in particular the rectal reporter array *gaIs245[col-34p::his-24::mcherry; unc-119(+)]* (Kagias et al., 2012) were used to identify K, K’ and K.p cells’ nuclei. Of note, K is found on the left side, K’ on the right side of the worm, and K.p is posterior to K.a and K’. We further confirmed the left position of K relative to the localisation of the commissures of the GABAergic neurons visualised with *oxIs12 [unc-47p::GFP]* and DNA replication was visualised using *gaIs245[col-34p::his-24::mcherry; unc-119(+)]*. The assessment of K division in *lin-5* mutant was performed by using *gaIs245* together with a fluorescent marker of the plasma membrane of rectal cells (*fpIs101[col-34p::ph::gfp; odr-1p::dsRed]*). In *egl-5(n945)* mutant, since *col-34* expression is absent in rectal cells, an *egl-5* reporter *(bxIs7[egl-5(6*.*5kb)::gfp; lin-15(+)])* was used to identify K.a and K.p. This cytoplasmic reporter did not allow us to estimate the orientation of K division based on K.a and K.p nuclei alignment or to measure their nuclear volumes. Similarly, the combination of markers that we used to simultaneously assess K division and DVB formation (*gaIs245* and *wyIs75* for *sox-2*), as well as the overall perturbed rectal area, precluded the quantification of the orientation of K division or the nuclear volumes in *sox-2* knockdown and *ceh-6* mutant. Since the rectal-specific *sox-2* antisense array contains also *rol-6(su1006)*, making it difficult to identify K.a, K’ and K.p in roller mutants, K cell division was scored in animals where *sox-2 is* knocked down by an anti-GFP nanobody (Wang et al., 2017) strategy (See RNAi/Silencing experiments). This strategy was also used in combination with *wrm-1* mutant to test the genetic interaction between *sox-2* and the Wnt signalling pathway. DVB presence was always based on the expression of *unc-47* terminal differentiation gene (*oxIs12, krIs6* or *wyIs75* arrays) and the presence of its stereotyped neurite going antero-ventrally. For timing experiments, tightly synchronized worms were obtained via hatch-pulses: more than 100-200 eggs were picked on fresh plates seeded with OP50 and each hour newly hatched larvae were transferred on new plates. Alternatively, L1-L2 worms were picked and staged according to the number of cells in the developing gonad. To determine the precise timing of K division and DVB formation, hatch-pulse tightly synchronized L1 were mounted every hour starting 10h after hatching. Several criteria were used to precisely stage the worms: the occurrence of division, using *gaIs245*; the L1-L2 transition, using the number of cells in the gonad and the disappearance of the alae observed by DIC. The early L2 stage was further dissected using the number of GABAergic cell bodies in the ventral nerve cord and the number of VD commissures as well as *unc-47* expression in DVB as observed with *oxIs12*.

### Confocal Microscopy

For confocal microscopy, worms were immobilized on 5% agarose pads with Tricaine 0.4% and Tetramizole 0.04% and captured on an inverted Leica TCS SP5 laser scanning confocal microscope (Leica Microsystems, Germany).

### Electron Microscopy

See description in Jarriault et al. 2008 (SI text). Ten microns of serial ultrathin sections (50-70 nm) of the rectal area were collected and contrasted in lead citrate and uranyl acetate before imaging with a SiS Megaview 3 CCD camera mounted on a FEI Morgagni TEM operated at 70 kV.

### Image processing and analysis

Images were processed using ImageJ. The measurement of the angle of K division was performed using the angle tool from ImageJ where a segmented line was drawn through the rectal slit and the center of the K.a and K.p nuclei in late L1 larvae as represented by the yellow dashed line in Figure 2A.

### Transcription factor binding sites analysis

For the manual identification of the putative TF binding sites, the following consensus sequences in Snapgene were used:

POP-1/TCF: WWCAAAR (Bertrand, 2009): SEM-4/SPALT: WARATTGTSTKKSW (Lloret-Fernandez, 2018) and TTGTST (Toker, 2003), SOX-2/SOX2: VACAAWGG (MAO143.3, JASPAR, Maruyama, 2005)

For automatic identification of putative binding sites, several platforms were used: FIMO with the *C*.*elegans* POP-1/TCF Matrix (Narasimhan, 2015), or MatInspector and Promo from vertebrate homologs.

Conservation of the binding sites was examined using the multiple alignment provided by the UCSC genome Browser. However, very poor conservation of the binding sites was observed among *C. elegans* species.

### 6×His-tagged protein expression and purification

To make 6×His-tagged proteins, *sox-2, pop-1* and *HMG-pop-1* cDNAs were cloned into pET32a+ or pET30a+ (Addgene) and transformed into BL21(DE3) from Stratagene. 6×His-tagged protein expression was to induced by adding 1mM IPTG to 1L of transformed cell culture at OD600 = 0.6 for 4 h at 37°C. Cells were then harvested and lysed in 40mL of Lysis Buffer (20 mM Tris pH 8, 100 mM NaCl, 10 mM imidazole, 1 mM DTT, 0.1 % NP40, Protease inhibitor from Roche) by sonication. The lysates were centrifuged 30 min at 40 000 RPM, 4°C. 6×His-tagged proteins were purified using a Hitrap Ni crude 1mL columns (Dutscher) and the following buffers: Equilibration Buffer (20 mM Tris pH 8, 100 mM NaCl, 10 mM imidazole, 0.1 % NP40) and Elution Buffer (Equilibration Buffer + 300 mM imidazole). The extracts were fractionated on a cation exchange chromatography and then dialyzed in EMSA buffer (20 mM Tris pH 8, 100 mM NaCl, 1 mM EDTA, 1 mM DTT, 0.1 % NP40). Note that for unknown reasons, we didn’t manage to purify the full length POP-1 protein despite numerous attempts.

### Electrophoretic mobility shift assay (EMSA)

Cy5-labeled probes were ordered (Merck, Germany) and annealed as follow: 10µM of forward (CY5) and reverse (unlabeled) primers were incubated in the dark in TE buffer containing 125mM NaCl at 100°C for 5min followed by 5hrs at room temperature and then O/N at 4°C. Purified 6×His-tagged proteins (250mM, or a range of 0,5 to 500nM) and 200ng of Cy5-probes were incubated on ice in binding buffer (20 mM Tris HCl pH 8.0, 100 mM NaCl, 100 µg/mL BSA, 1 mM EDTA, 1 mM DTT, 0.1% NP-40) for 20 min and resolved in a pre-run 6% polyacrylamide gel containing 0.5× TBE buffer (Bio-Rad Mini-PROTEAN) at 100 V for 1 h. The gels were imaged using the Typhoon™ FLA 9500 biomolecular imager.

## QUANTIFICATION AND STATISTICAL ANALYSIS

### Statistical analysis and data representations

In the histograms, mean and standard deviation between biological replicates of the percentage of worms scored are represented. The stars summarize the statistical significance as calculated through Fisher’s exact test on the merged raw data from single replicates in a contingency table. Two-tailed Fisher’s exact test was used to compare reporter gene expression in mutant vs wild type worms. One-tailed Fisher’s test was used to compare Td defects in mutant vs wild type worms; the choice of the one-tailed test is justified by the known 0% Td defect in wild type worms. The Student’s t-test was used to analyze the significance in the difference between K.a and K.p nuclear volumes’ ratio and the angle of K division in wild type vs *lin-17* and *sem-4* mutants. F test was used to compare the variances of the angle of division between wild type vs *lin-17, sem-4* and *goa-1* mutants. * p<0.05; ** p<0.01; *** p<0.001; **** p<0.0001 and ns, not significant, for all the tests.

### Quantification of nuclear volumes

To measure K.a, K.p and K’ nuclear volumes, *gaIs245* transgenic strains were imaged at Leica SP5 confocal microscope. All the volume in Z containing K.a, K.p and K’ nuclei was acquired with a z-step size of 0.3 µm. Imaris software was used to reconstruct 3D images and to analyze the volume occupied by each nuclei. Manual selection of the nuclear area was performed.

## Supplementary figure legends

Statistical tests, see Star Methods, unless otherwise stated. In all figures., n, total animal scored.

**Supplementary Fig. 1. K replication is not sufficient for DVB formation and G**.

(A) Histograms summarizing the percentage of animals in which K DNA underwent replication in the *lin-5(ev571ts)* mutant at different restrictive temperatures.

(B) Histograms showing the percentage of worms without DVB in mutants for the Gα genes and *gpr-1/LGN* involved in spindle orientation in *C. elegans* zygote and the Par gene *par-1*. The low penetrance of DVB absence is due to an impairment in K cytokinesis.

(C) Dot plot representing K division angle in the *goa-1(sa734)* mutant. n=64. ns, not significant.

**Supplementary Fig. 2. Time course of K division and DVB formation in the wild type background**.

(A) Time course of the cellular events during K division and DVB formation along the developmental timeline. The *col-34p::mCherry* positive rectal cells and the rectal slit (line) are represented. Each box corresponds to a point of one hour where different characteristic landmarks have been observed in addition to K division: number of GABAergic neurons, VD commissures, presence of alae, number of cells in the gonad, *col-34p::mCherry* and *unc-47p::GFP* intensities, rectal cell shapes. The percentage of animals (n%) with the corresponding landmarks is indicated at each particular time point. Worms were synchronized by hatch pulse (see Methods). The left side of the animal is towards front, anterior to the left, and dorsal is up.

(B) Quantification of worms with expression of *fpEx1062[let-413::gfp::pest*], *fpIs110[lin-26p::GFP]* and *oxIs12[unc-47p::GFP]* in K.p over time, in L1 and L2 grown at 20°C.

**Supplementary Fig. 3. The non-canonical Wnt pathways are not required for K-to-DVB**.

(A) Histograms showing the percentage of “No DVB” worms in mutant backgrounds for genes of the PCP pathway.

(B) Histograms showing the percentage of “No DVB” worms in mutant backgrounds for the non-canonical Wnt-dependent pathways (*lin-18* and *cam-1*) or their downstream effectors (*ced-10*). ns, non-statistically significant; **, p<0.005.

**Supplementary Fig. 4. The K.p cell remains rectal-epithelial in *lin-17/FZD* and *sem-4/SALL* mutants**.

Quantification of the % of animals expressing (A-C) Epithelial (*ajm-1, let-413* and *lin-26*), (D) rectal (*egl-5*), (E-G) pan-neuronal (*unc-119, unc-33* and *rgef-1*) and (H, I) GABAergic (*unc-25, unc-47*) reporters in K posterior daughter in *lin-17/F*ZD and *sem-4/SALL* mutant backgrounds, or DVB in wild type, in L4 larvae.

**Supplementary Fig. 5. The K.p cell expresses the apical junction protein AJM-1 in *lin-17/FZD* and *sem-4/SALL* mutants**.

Confocal images of wild type, *lin-17/F*ZD and *sem-4/SALL* mutant background in L3 larvae carrying *gaIs245[col-34p::his-24::mcherry]* to visualize the rectal cell nuclei and *jcIs1[ajm-1::GFP]*. Patches of AJM-1 protein are present in the K.p cell (dashed oval) in the mutant background.

**Supplementary Fig. 6. *sox-2, ceh-6* and *egl-27* paralogs do not seem to be required to form DVB**.

Quantification of DVB defective L4 animals (as observed by *unc-47* expression) using RNAi in a sensitized *rrf-*3 mutant background to target *sox-2* paralogs (*dpy-8* and *gfp* RNAi represent controls). Mutants were used for paralogs of *ceh-6* (*unc-86(n846)* and *ceh-18(mg57)*) and *egl-27* (*lin-40(ku285)*). No obvious defects were observed, although RNAi was found to work poorly in the rectal cells.

**Supplementary Fig. 7. *sem-4/SALL* and the Wnt signaling pathway act in parallel to drive K-to-DVB Td**.

(A) Quantification of *sem-4/SALL* expression in K.p in wild type L4s and in *lin-17/Frizzled* mutant L4s.

(B) Quantification of *lin-17/FZD* expression in K.p in wild type L4s *vs sem-4/SALL* and *lin-17/FZD* mutants.

(C-D) Quantification of DVB defective L4 animals (as observed by *unc-47* expression using *krIs6* in A and *oxIs12* in B) in simple *sem-4(n1378)* (C), *sox-2* knock-down (using a nanobody strategy, D) and *wrm-1(n1982)* (C, D) mutants, or in *sem-4(n1378);wrm-1(n1982)* (C) and *wrm-1(n1982);sox-2* KD (D) double mutants, all raised at 25°C.

**Supplementary Fig. 8. *lim-6* intron 4 binding sites analysis**.

(A) The intron 4 of *lim-6* was analyzed using different tools (See Methods). Here are represented the POP-1 binding site prediction using the Matrix of Narasimhan and the consensus of Bertrand, 2009. The SOX-2 binding sites were predicted by Promo and the SEM-4 consensus site published in Toker, 2003 was used.

(B) Sequence logo for POP-1 and SOX-2 binding sites showing the sequence similarities.

(C) Table summarizing the binding sites on which we have focused our efforts in this study. Binding sites predicted by more than one approach, or because of the presence of two consecutive binding sites for SOX-2 (site 4), were selected. The mutations introduced into the *lim-6* transcriptional reporter to abolish POP-1 binding are presented on the right. Note that these mutations most probably abolish also SOX-2 binding due to the very close similarity of their predicted binding sites. (D) This binding site similarity can be explained by the sequence similarity of the HMG domains present in SOX-2 and POP-1.

(E) Sequence of the probes used in this study for the gel shift experiments.

**Supplementary Fig. 9. *sox-2* and *lim-6* expression in K posterior daughter**.

(A) Fluorescent images of *gfp::sox-2* CRISPR (KI) and *col-34p::his-24::mcherry* in the rectum of an L1 animal before the division (top), in an L1 animal after the division (14 cells in the gonad; middle) and in an L2 animal (bottom). Note that K.a continues to express *sox-2* over time whereas expression fades away in K.p during its conversion. White stars indicate the rectal gland cells; dashed line, rectal slit; anterior is to the left and dorsal up.

(B) Quantification of the % of L4 larvae expressing *lim-6::gfp* CRISPR (KI) and *lim-6 intron 4* transcriptional reporter (*int4*) in wild-type (DVB) and *pop-1(q645)* mutant (persistent K.p) backgrounds. Note that for the *pop-1(q645)* mutant, only viable homozygote (not balanced) mutant worms were analyzed.

**Supplementary Fig. 10. Gel shift experiments showing independent binding capacities of POP-1 and SOX-2 to probes 1, 2 and 4**.

Increasing concentration (5nm, 50nM and 500nM) of purified HMG-POP-1 and SOX-2 were incubated with wild type or mutated probes 1, 2 and 4 bound to the Cy5 fluorophore. Note that the probe 4 which does not bear canonical SOX-2 binding site is able to bind SOX-2.As, in addition, the mutation of the POP-1 binding site doesn’t seem to affect this binding, it is likely that a non-predicted SOX-2 binding site is present. Probe 3 was also able to bind both SOX-2 and HMG-POP-1, although results for are not presented because this probe annealed poorly, most probably due to its AT-rich sequence. Blue arrowhead, POP-1 bound to the probe; orange arrowhead, SOX-2 bound to the probe; light orange arrowhead, a second SOX-2 shifted band appears at high SOX-2 concentrations; open arrowhead, unbound probe.

**Supplementary Fig. 11. Gel shift experiments showing binding capacities of POP-1 and SOX-2 when co-incubated**.

(A) Increasing quantity of SOX-2 (125nM-250nM-500nM) was added to a mix of HMG-POP-1 and Cy-5-dsProbe #1, #2, *odr-1* (known SOX-2 target, Alqadah, 2015) and *ceh-22* (known POP-1 target, Lam, 2007, Bhambhani, 2014). Note that SOX-2-Cy-5-dsprobe control, as was presented in main Fig. 5, was not added in parallel here.

(B) Increasing quantity of HMG-POP-1 (125nM-250nM-500nM) was added to a mix of SOX-2 and Cy-5-dsProbe #1,# 2, #4 and *ceh-22*. Increasing quantity of HMG-POP-1 shows an increasing binding to all the probes as well as an upper shift, most probably corresponding to a HMG-POP-1-SOX-2-Probe complex.

Blue arrowhead, POP-1 bound to the probe; orange arrowhead, SOX-2 bound; green arrowhead, POP-1 and SOX-2 bound; open arrowhead, unbound probe.

