## Supplementary material for "A natural transdifferentiation event involving mitosis is empowered by integrating signaling inputs with conserved plasticity factors": Table S1

**Supplementary Table 1. List of reporters' expression.**

| *C. elegans* genes | Human ortholog | Reporter | WT L1 K | WT early L2 K.a | WT early L2 K.p | WT L4 DVB | *sem-4* L4 K post. daughter | *lin-17* L4 K post. daughter | References reporter | Ref observation |
|---|---|---|---|---|---|---|---|---|---|---|
| **Epithelial markers** | | | | | | | | | | |
| ***dlg-1*** | *DLG* | *mcIs46[dlg-1::rfp]* | + | + | - | - | N.D | N.D | Diogon et al. (2007) | This study |
| ***ajm-1*** | *AJM1* | *jcIs1[ajm-1::gfp]* | + | + | - | - | + | + | Mohler et al. (1998) | This study; Mohler et al. (1998) |
| ***hmr-1*** | Cadherin | *fpIs17[hmr-1::gfp]* | + | + | - | - | N.D | N.D | This study | This study |
| ***let-413*** | *SCRIB* | *fpEx1062[let-413a::gfp::pest]* | + | + | + | - | + | + | This study | This study |
| ***lin-26*** | Zinc-finger transcription factor | *fpIs110[lin-26rectalp::gfp]* | + | + | + | - | + | + | Labouesse et al. (1996); this study | Labouesse et al. (1996); this study |
| **Rectal markers (also in other cells, to visualise the rectal cells)** | | | | | | | | | | |
| ***sem-4*** | *SALL* transcription factor | *syb1287[sem-4::gfp]* | + | + | + | + | ND | + | This study | This study |
| ***sox-2*** | *SOX* transcription factor | *syb737[gfp::sox-2]* | + | + | +* | - | + | + | This study | This study |
| ***ceh-6*** | *POU* transcription factor | *syb972[gfp::ceh-6]* | + | + | +* | - | + | + | This study | This study |
| ***egl-5*** | *HOX* transcription factor | *bxIs7[egl-5::gfp]* | + | + | + | - | ND | ND | Teng et al. (2004) | This study |
| ***col-34*** | Cuticle collagen gene | *gaIs245[col-34p::his-24::mcherry]* | + | + | + | - | + | + | Zuryn et al. (2014) | This study |
| ***got-1.2*** | *GOT1* | *sIs11174[rCesT01C8.5::gfp+pCeh361]* | + | + | + | - | ND | ND | McKay et al. (2003) | This study |
| **Pan-neuronal markers** | | | | | | | | | | |
| ***unc-33*** | *DPYS* | *otIs117[unc-4(+) ; unc-33p::GFP]* *otIs118[unc-33::GFP; unc-4(+)]* | - | - | - | + | - | - | McKay et al. (2003) | This study |
| ***unc-119*** | *UNC119* | *edIs6[unc-119::gfp; rol-6]* | - | - | - | + | - | - | Maduro and Pilgrim (1995); Praitis et al. (2001) | This study |
| ***rgef-1*** | RASGRP3 | *otIs173 [F25B3.3::DsRed2; ttx-3promB::GFP]* | - | - | - | + | - | - | Benard et al. (2009) | This study |

| | | | | | | | | | | |
|---|---|---|---|---|---|---|---|---|---|---|
| **DVB Terminal selector** | | | | | | | | | | |
| ***lim-6*** | *LMX1B* | *syb971[lim-6::gfp]* | - | - | + | + | - | - | This study | Hobert et al. (1999); this study |
| **GABAergic markers** | | | | | | | | | | |
| ***unc-47*** | *SLC32A1* | *oxIs12[unc-47p::gfp]* | - | - | - | + | - | - | McIntire et al. (1997) | McIntire et al. (1997); this study |
| | | *krIs6[unc-47p::DsRed2]* | | | | | | | Teuliere et al. (2011) | Teuliere et al. (2011) |
| ***unc-25*** | *GAD* | *juIs8[unc-25p::gfp]* | - | - | - | + | - | - | Jin et al. (1999) | Jin et al. (1999) ; this study |

*, expression is seen in K.p after its birth, and disappears as *lim-6* expression appears (see Fig. 5A).


Benard, C., Tjoe, N., Boulin, T., Recio, J., and Hobert, O. (2009). The small, secreted immunoglobulin protein ZIG-3 maintains axon position in Caenorhabditis elegans. Genetics *183*, 917-927.

Diogon, M., Wissler, F., Quintin, S., Nagamatsu, Y., Sookhareea, S., Landmann, F., Hutter, H., Vitale, N., and Labouesse, M. (2007). The RhoGAP RGA-2 and LET-502/ROCK achieve a balance of actomyosin-dependent forces in C. elegans epidermis to control morphogenesis. Development *134*, 2469-2479.

Hobert, O., Tessmar, K., and Ruvkun, G. (1999). The Caenorhabditis elegans lim-6 LIM homeobox gene regulates neurite outgrowth and function of particular GABAergic neurons. Development *126*, 1547-1562.

Jin, Y., Jorgensen, E., Hartwieg, E., and Horvitz, H.R. (1999). The Caenorhabditis elegans gene unc-25 encodes glutamic acid decarboxylase and is required for synaptic transmission but not synaptic development. J Neurosci *19*, 539-548.

Labouesse, M., Hartwieg, E., and Horvitz, H.R. (1996). The Caenorhabditis elegans LIN-26 protein is required to specify and/or maintain all non-neuronal ectodermal cell fates. Development *122*, 2579-2588.

Maduro, M., and Pilgrim, D. (1995). Identification and cloning of unc-119, a gene expressed in the Caenorhabditis elegans nervous system. Genetics *141*, 977-988.

McIntire, S.L., Reimer, R.J., Schuske, K., Edwards, R.H., and Jorgensen, E.M. (1997). Identification and characterization of the vesicular GABA transporter. Nature *389*, 870-876.

McKay, S.J., Johnsen, R., Khattra, J., Asano, J., Baillie, D.L., Chan, S., Dube, N., Fang, L., Goszczynski, B., Ha, E.*, et al.* (2003). Gene expression profiling of cells, tissues, and developmental stages of the nematode C. elegans. Cold Spring Harb Symp Quant Biol *68*, 159-169.

Mohler, W.A., Simske, J.S., Williams-Masson, E.M., Hardin, J.D., and White, J.G. (1998). Dynamics and ultrastructure of developmental cell fusions in the Caenorhabditis elegans hypodermis. Curr Biol *8*, 1087-1090.

Praitis, V., Casey, E., Collar, D., and Austin, J. (2001). Creation of low-copy integrated transgenic lines in Caenorhabditis elegans. Genetics *157*, 1217-1226.
Teng, Y., Girard, L., Ferreira, H.B., Sternberg, P.W., and Emmons, S.W. (2004). Dissection of cis-regulatory elements in the C. elegans Hox gene egl-5 promoter. Dev Biol *276*, 476-492.

Teuliere, J., Gally, C., Garriga, G., Labouesse, M., and Georges-Labouesse, E. (2011). MIG-15 and ERM-1 promote growth cone directional migration in parallel to UNC-116 and WVE-1. Development *138*, 4475-4485.

Zuryn, S., Ahier, A., Portoso, M., White, E.R., Morin, M.C., Margueron, R., and Jarriault, S. (2014). Transdifferentiation. Sequential histone-modifying activities determine the robustness of transdifferentiation. Science *345*, 826-829.
