## Supplementary figures and images for "A natural transdifferentiation event involving mitosis is empowered by integrating signaling inputs with conserved plasticity factors"

### Fig. S1

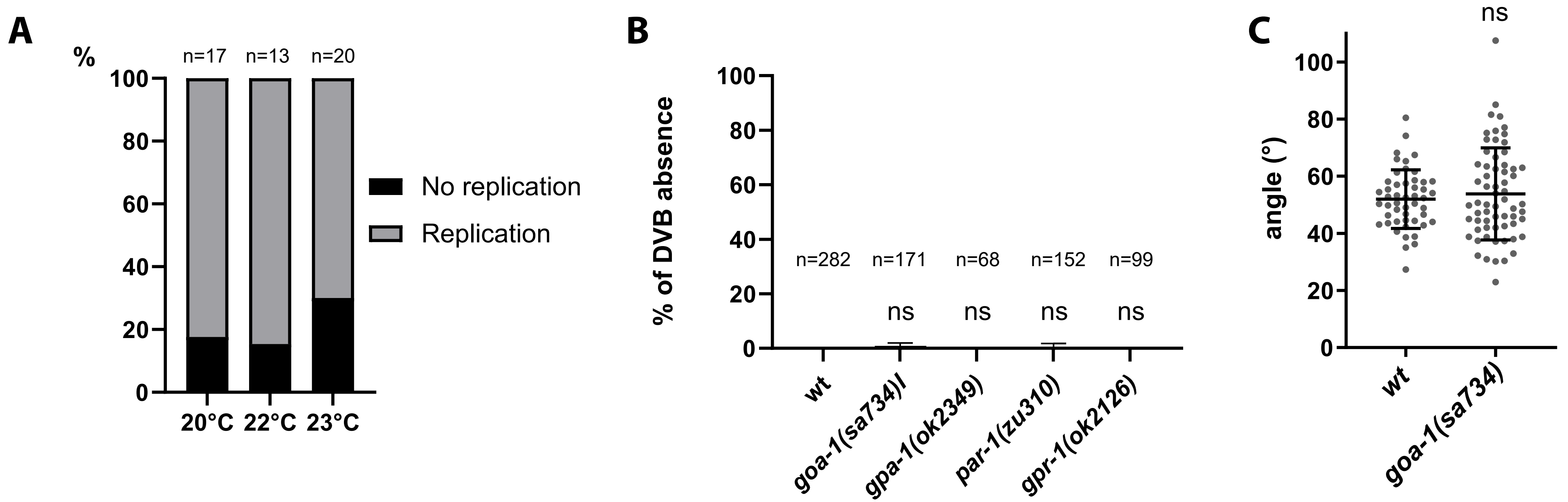


**Supplementary Figure 1**

### Fig. S2

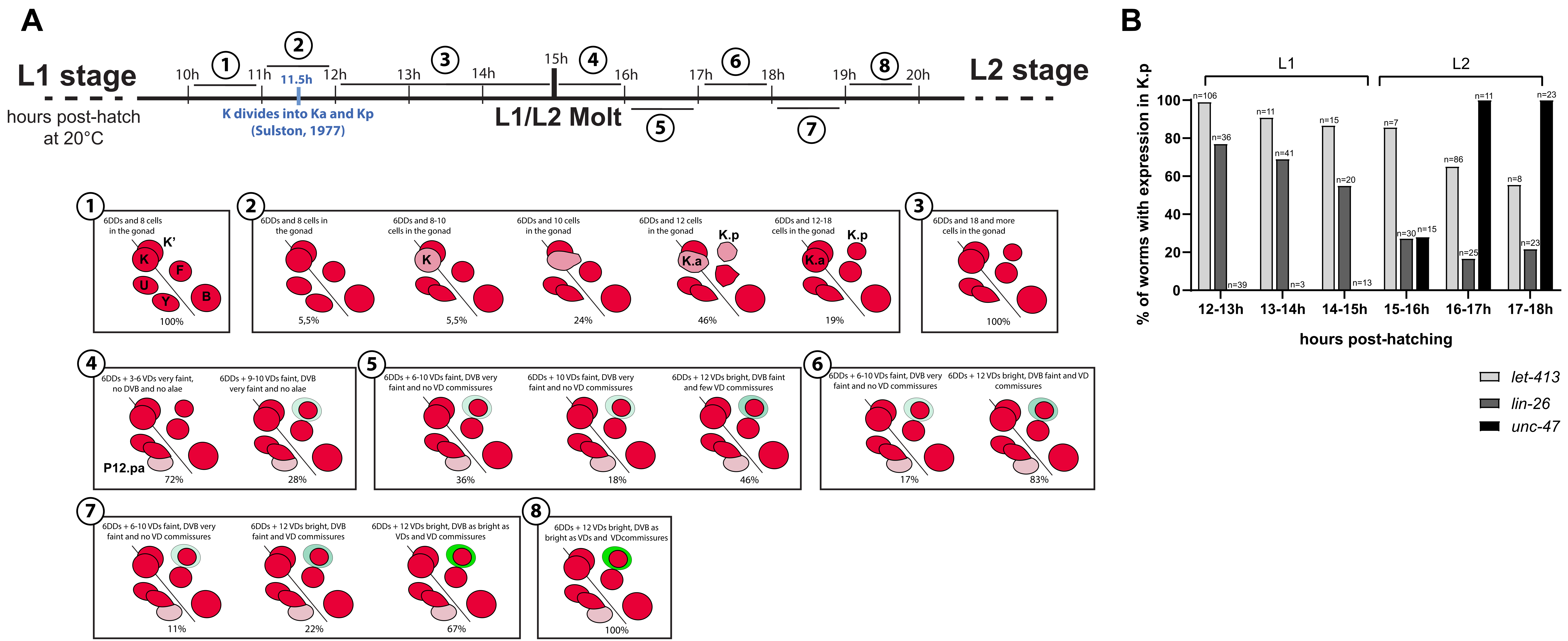

### Fig. S3

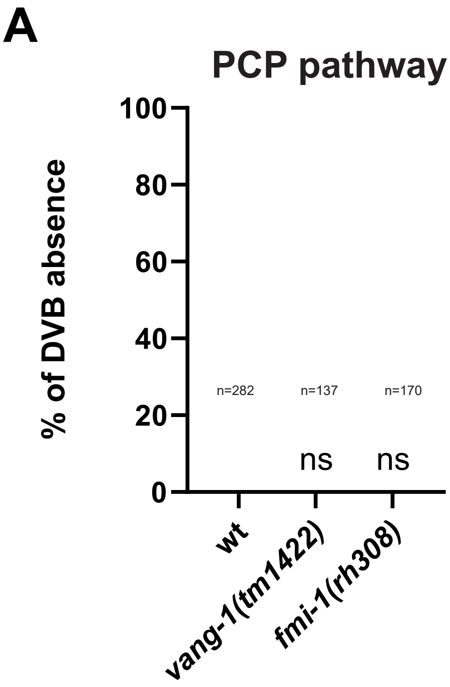


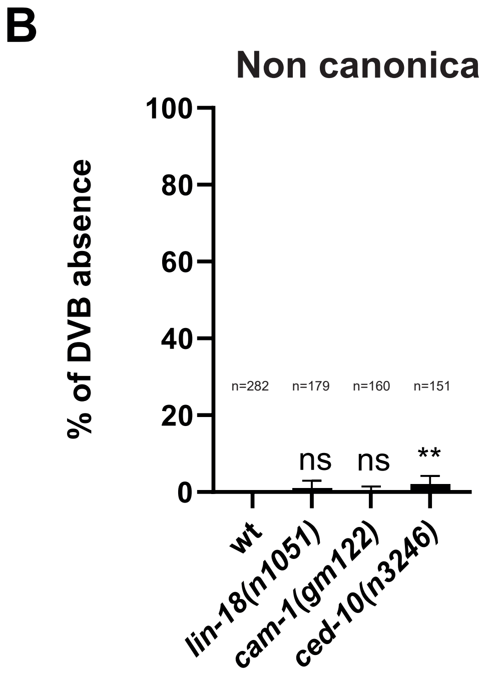


Supplementary Figure 3

### Fig. S4

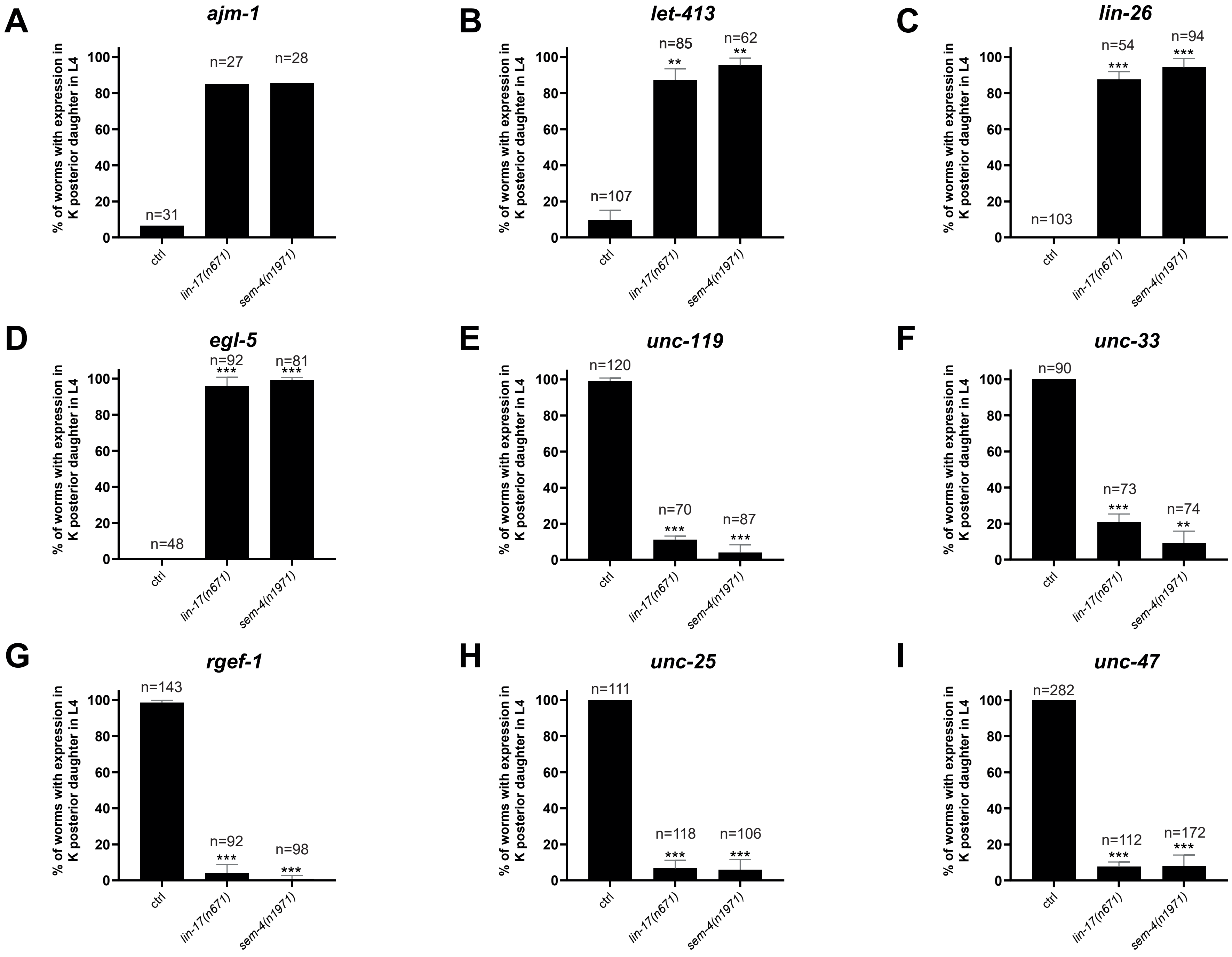


**Supplementary Figure 4**

### Fig. S5

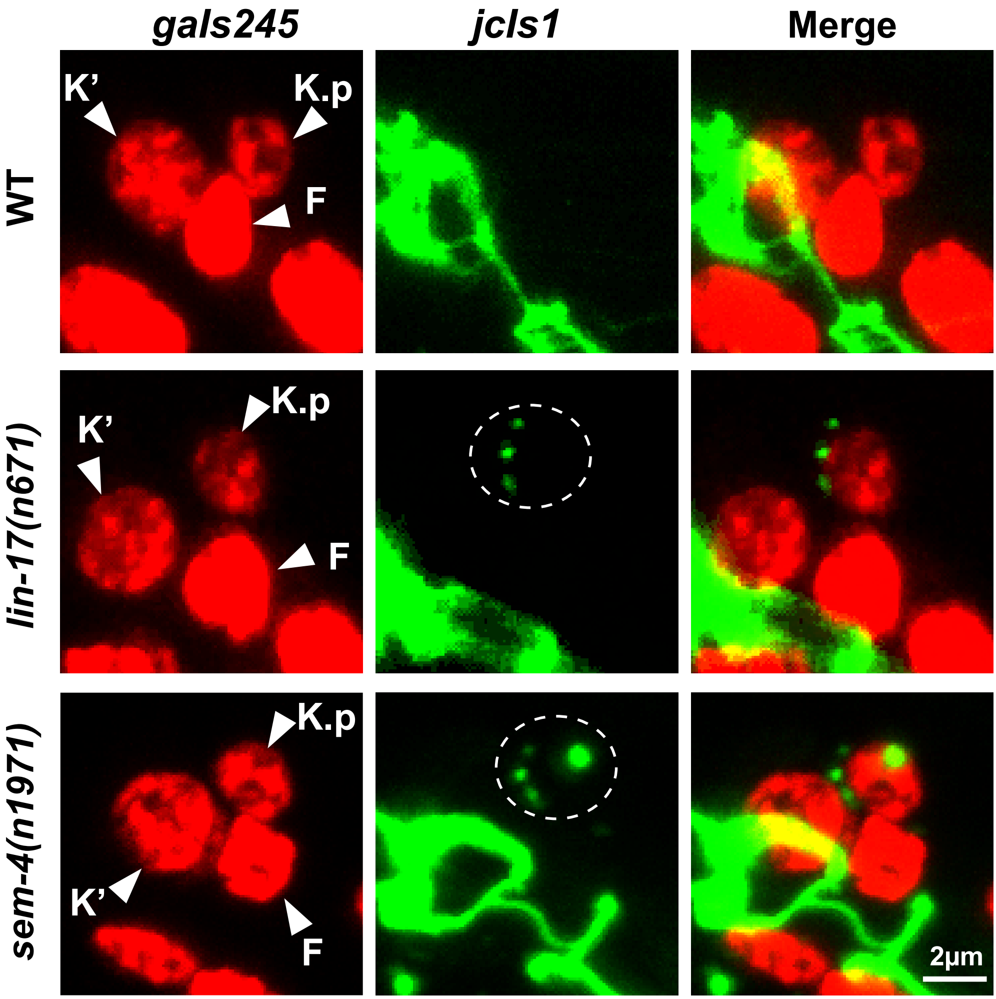


**Supplementary Figure 5**

### Fig. S6

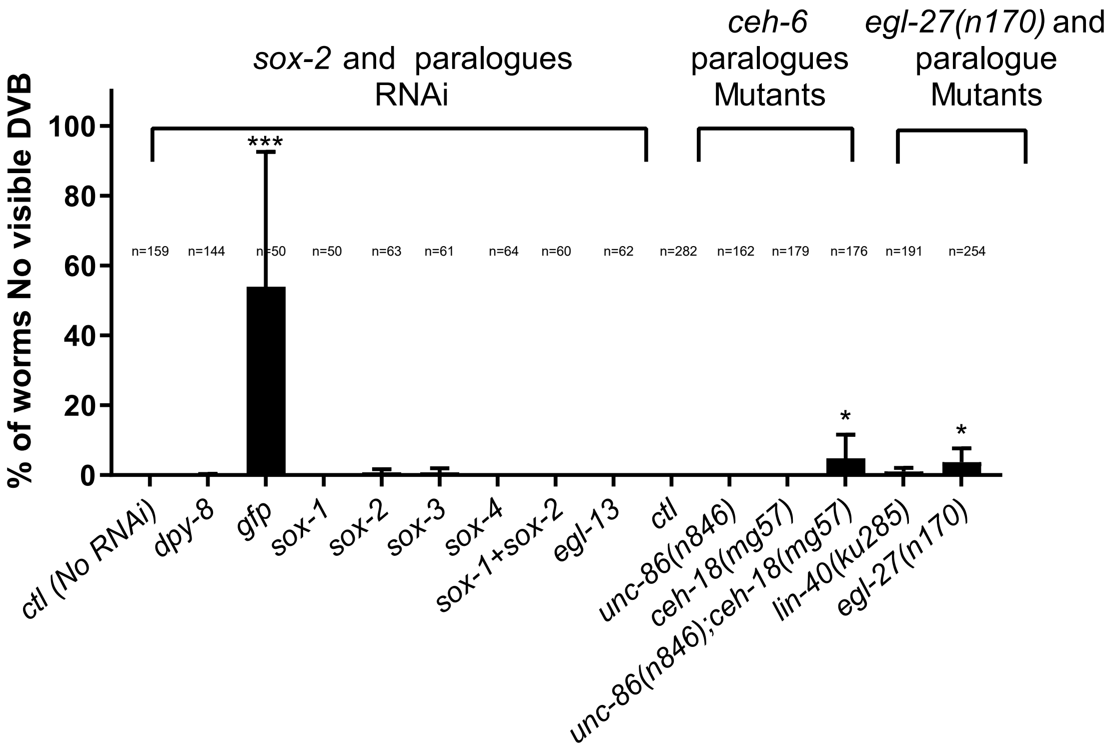


**Supplementary Figure 6**

### Fig. S7

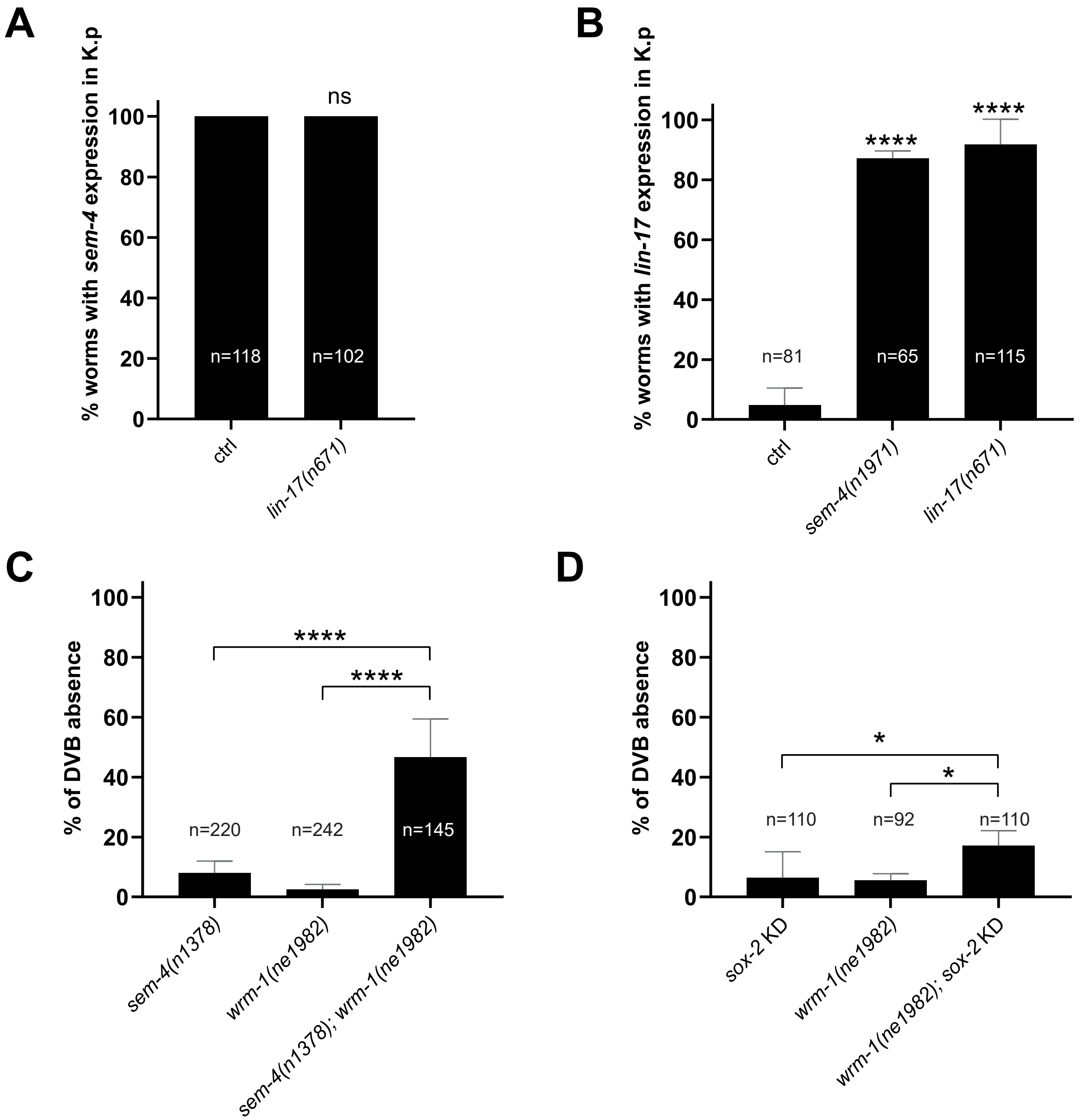

### Fig. S9

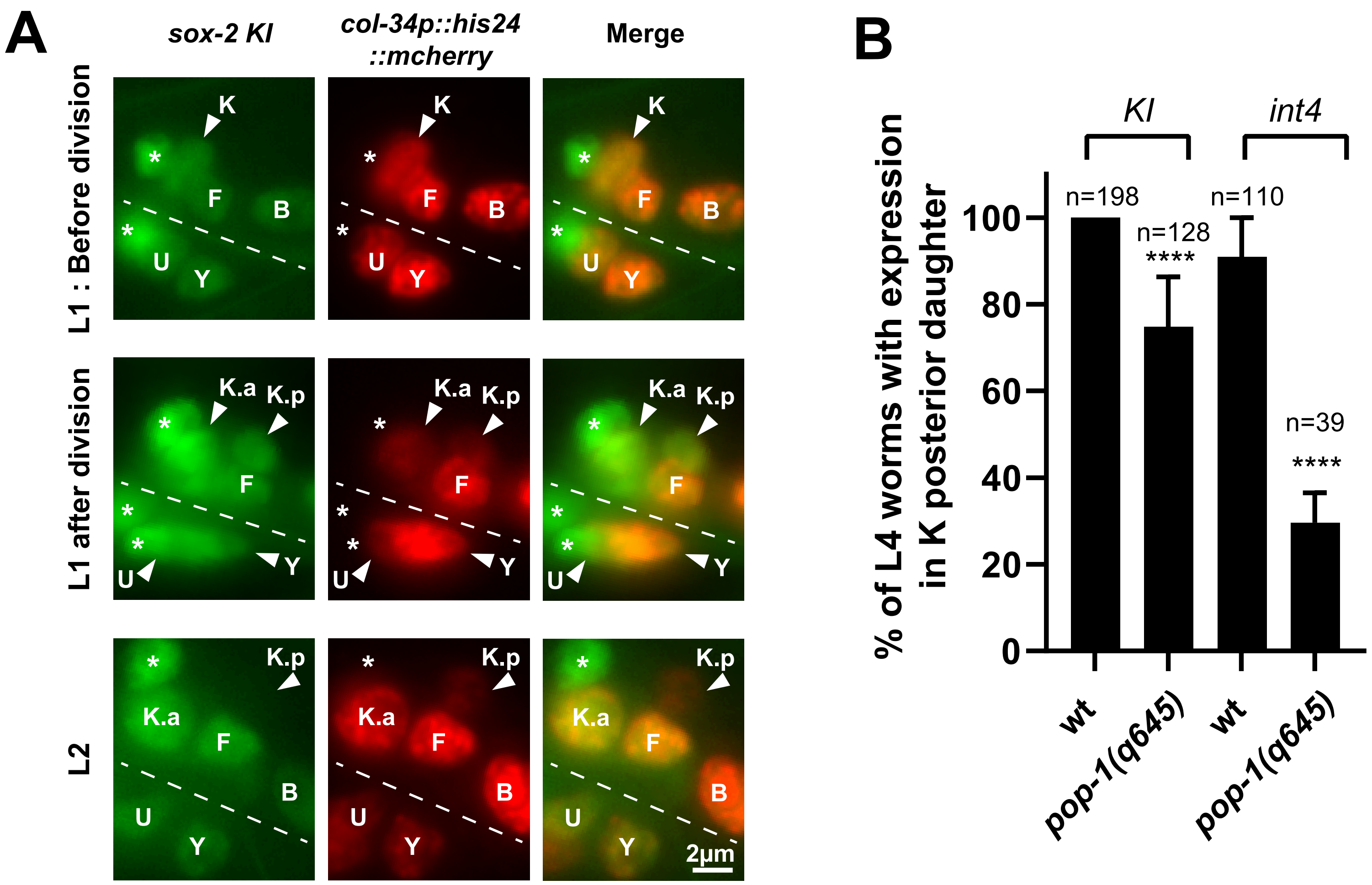

### Fig. S10

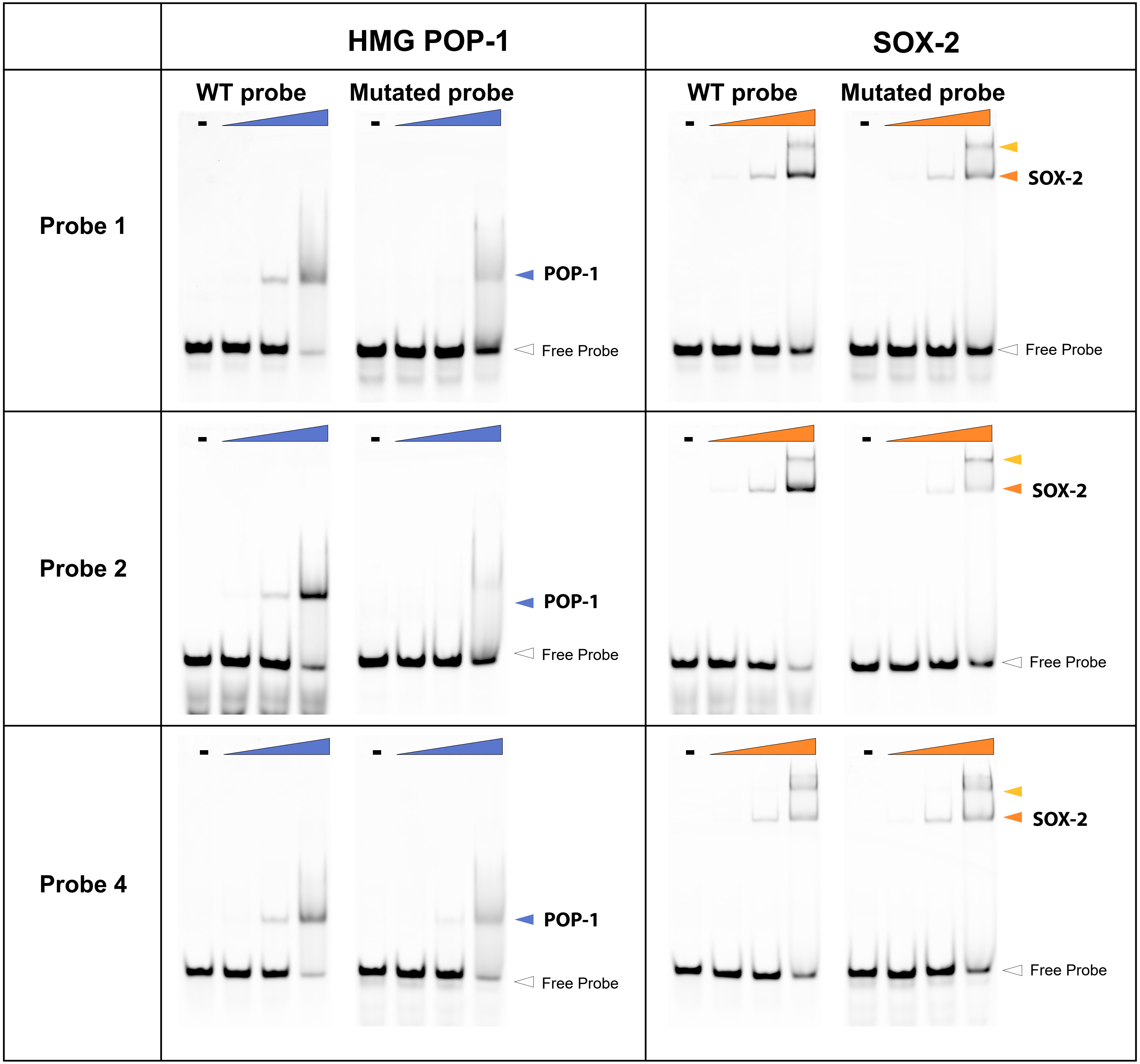


Sup Figure 10

### Fig. S11

Sup Figure 11

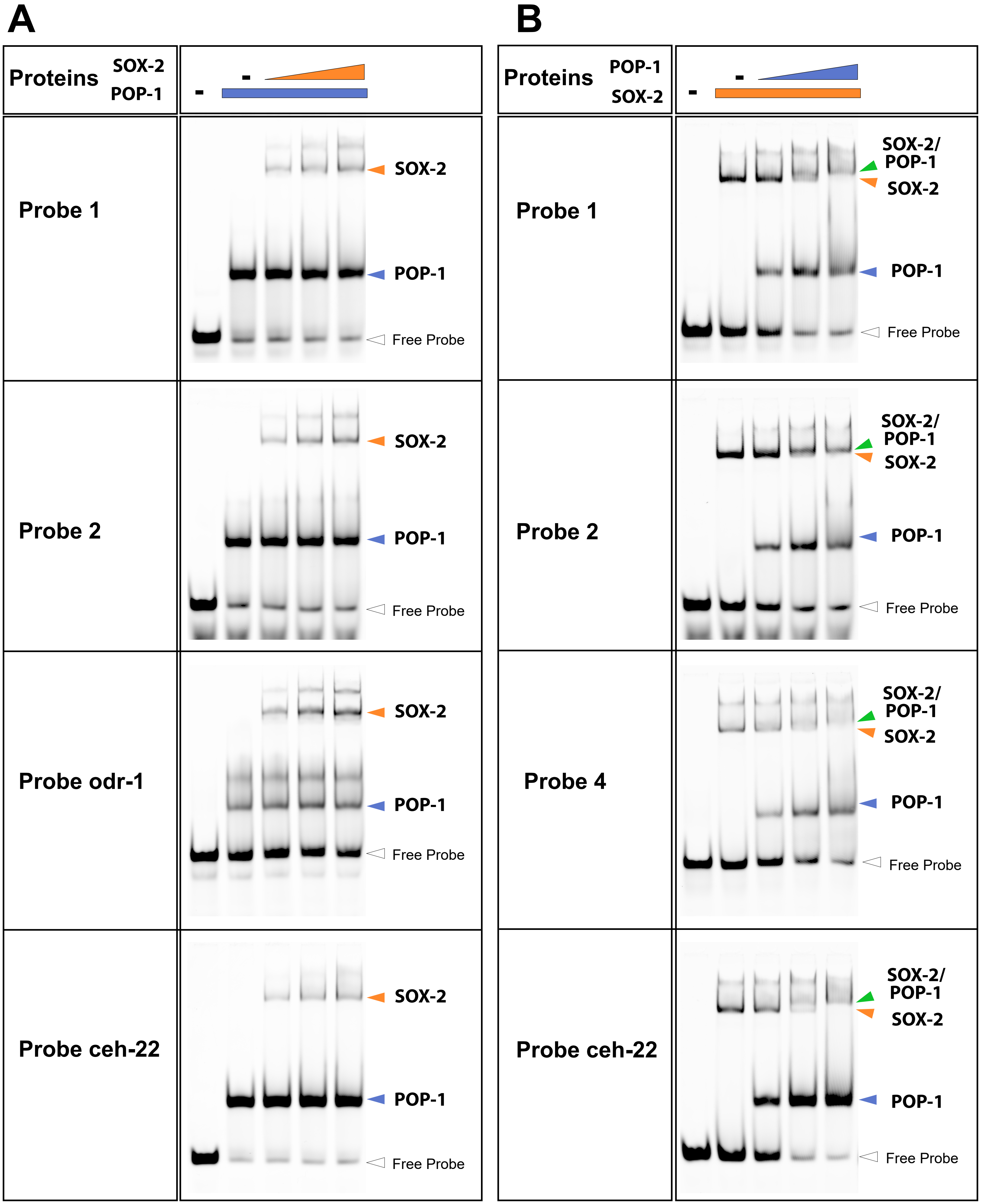
