## Supplementary material for "A natural transdifferentiation event involving mitosis is empowered by integrating signaling inputs with conserved plasticity factors": Fig. S8

### Sup Figure 8

**A**

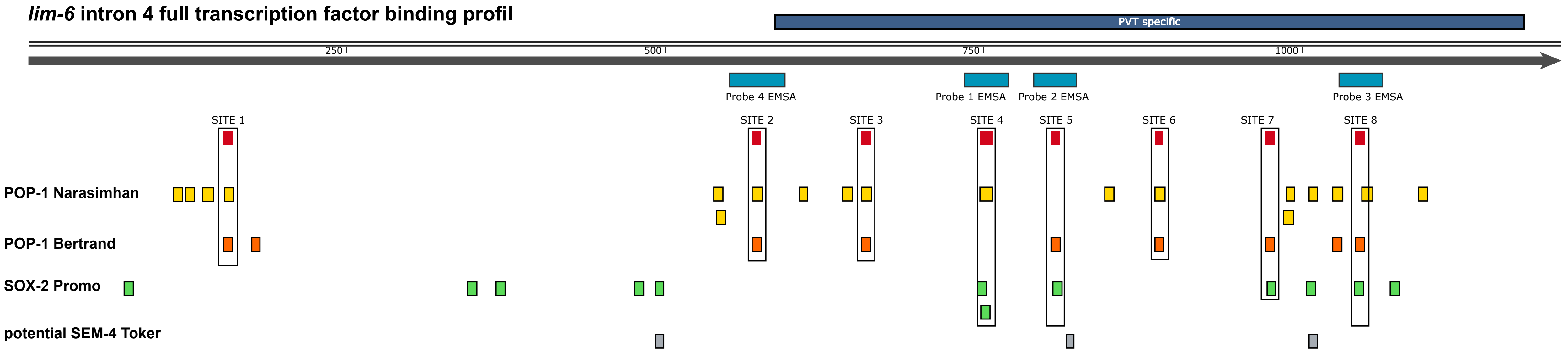


**B**

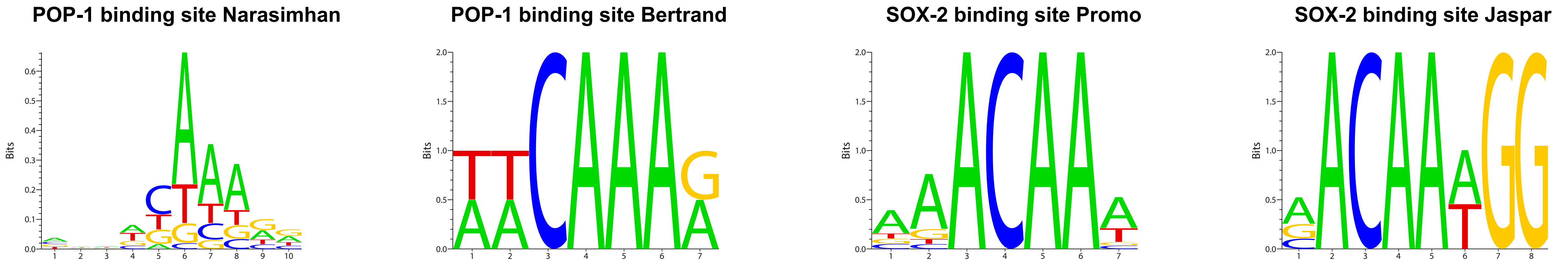


**C**

| Site | Sequence | 1st nucleotide Position in intron 4 (bp) | Identification method | Mutation in pSJ759 |
|---|---|---|---|---|
| **Site 1** | TTCAAAG | 154 | Fimo: Narasimhan, 2015<br>Bertrand, 2009<br>MatInspector | |
| **Site 2** | ATCAAAA | 569 | Fimo: Narasimhan, 2015<br>Bertrand, 2009<br>MatInspector | ATCGTTA |
| **Site 3** | TTCAAAG | 655 | Fimo: Narasimhan, 2015<br>Bertrand, 2009<br>MatInspector | TTCGTTG |
| **Site 4** | ACAACAAACA | 748 | Fimo: Narasimhan, 2015 | ACACCATGGA |
| **Site 5** | CTTTGTT | 804 (rev) | Fimo: Narasimhan, 2015<br>Bertrand, 2009<br>MatInspector | CGCTAGC |
| **Site 6** | ATCAAAG | 885 | Fimo: Narasimhan, 2015<br>Bertrand, 2009<br>MatInspector | ATCCTCG |
| **Site 7** | CTTTGTT | 972 (rev) | Bertrand, 2009<br>MatInspector | CCCGCGG |
| **Site 8** | AACAAAA | 1043 | Bertrand, 2009<br>MatInspector | CGGCCGA |

**D**

**HMG**

```
SOX-2    RVKRPMNAFMVWSRGQRKKMA--LENPKMHNSEISKRLGTEWKMLSEQEKRPFIDEAKRLRAIHMKEHPDYKY   71
POP-1    HVKKPLNAFMWFMKENRKALLEEIGNNEKQSAELNKELGKRWHDLSKEEQAKYFEMAKKDKETHKERYPEWSA   73
         :**:*:**** : : :** :   : * : :.:*:.*.**..*: **::*:  ::: **: :  * :.:*::.
```

**E**

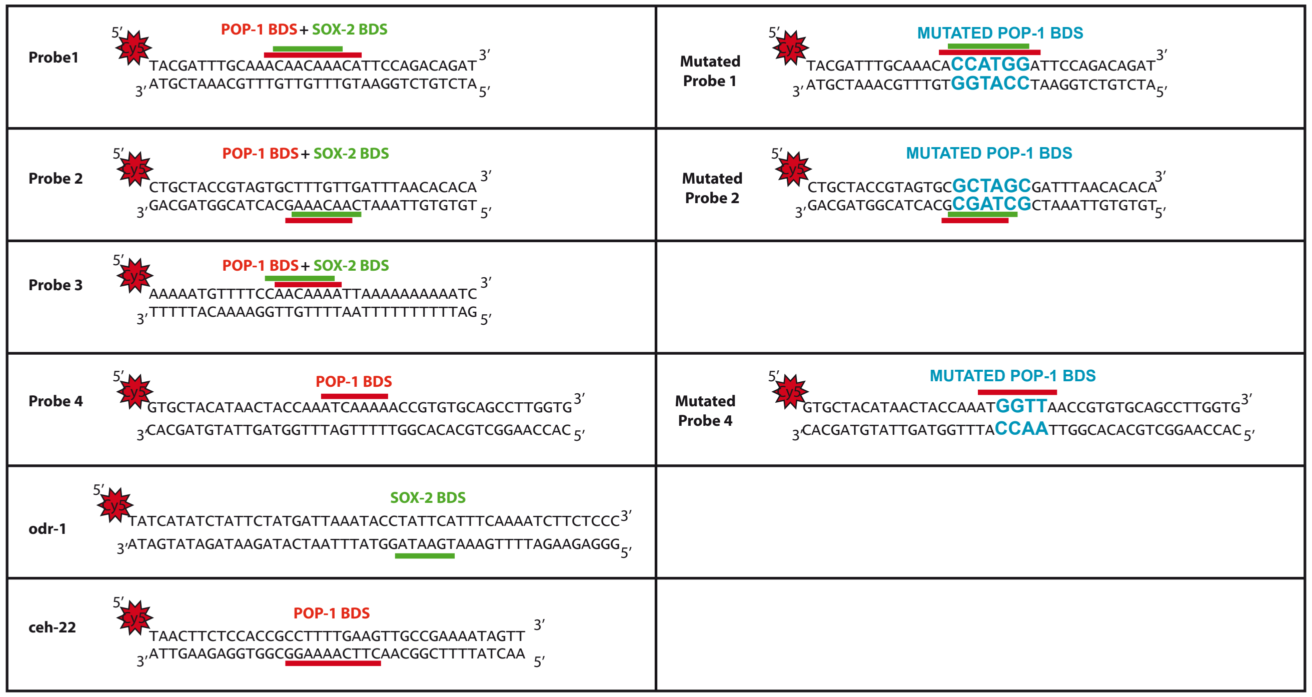
