## Supplementary material for "A natural transdifferentiation event involving mitosis is empowered by integrating signaling inputs with conserved plasticity factors": Table S2

### Supplementary Table 2. Strain list.

| C. elegans strain | Identifier |
| --- | --- |
| *rrf-3(pk1426) II ; oxIs12[unc-47::gfp; lin-15(+)] X* | IS17 |
| *lin-5(ev571) II; gaIs245[col-34p::his-24::mcherry; unc-119(+)] V ; oxIs12[unc-47p::gfp; lin-15(+)] X* | IS1118 |
| *sem-4(n1971) bxIs7[egl-5p(6,5kb)::gfp; lin-15(+)] I; otIs173[rgef-1p::dsred2; ttx-3p::gfp] III* | IS1208 |
| *sem-4(n197) I; otIs117[unc-4(+); unc-33p::gfp] IV; gaIs245[col-34p::his-24::mcherry; unc-119(+)] V* | IS1210 |
| *gaIs245[col-34p::his-24::mcherry; unc-119(+)] V; oxIs12[unc-47p::gfp; lin-15(+)] X* | IS1299 |
| *egl-5(n945) III; gaIs245[col-34p::his-24::mcherry; unc-119(+)] V; oxIs12[unc-47p::gfp; lin-15(+)] X* | IS1332 |
| *lin-17(n671) I; gaIs245[col-34p::his-24::mcherry; unc-119(+)] V; oxIs12[unc-47p::gfp; lin-15(+)] X* | IS1370 |
| *fpIs17[hmr-1::gfp]; gaIs245[col-34p::his-24::mcherry; unc-119(+)] V* | IS1374 |
| *wrm-1(ne1982) III; gaIs245[col-34p::his-24::mcherry; unc-119(+)] V; oxIs12[unc-47p::gfp; lin-15(+)] X* | IS1432 |
| *sem-4(n1971) I; gaIs245[col-34p::his-24::mcherry; unc-119(+)] V; oxIs12[unc-47p::gfp; lin-15(+)] X* | IS2968 |
| *unc-86(n846) III; gaIs245[col-34p::his-24::mcherry; unc-119(+)] V; oxIs12[unc-47p::gfp; lin-15(+)] X* | IS3097 |
| *fpIs110[lin-26p::gfp; rol-6(su1006)] IV; gaIs245[col-34p::his-24::mcherry; unc-119(+)] V* | IS3107 |
| *egl-27(ok1670) II; gaIs245[col-34p::his-24::mcherry; unc-119(+)] V ; oxIs12[unc-47p::gfp; lin-15(+)] X* | IS3113 |
| *gaIs245[col-34p::his-24::mcherry; unc-119(+)] V; fpEx1062[let-413a::gfp::pest; myo-2p::gfp]* | IS3119 |
| *gaIs245[col-34p::his-24::mcherry; unc-119(+)] V; oxIs12[unc-47p::gfp; lin-15(+)] X; fpEx955[Δ(-2846pb to -102)ceh-6p::gfp::ceh-6; odr-1::rfp]* | IS3120 |
| *ceh-6(gk665) I; gaIs245[col-34p::his-24::mcherry; unc-119(+)] V; oxIs12[unc-47p::gfp; lin-15(+)] X; fpEx955[Δ(-2846pb to -102)ceh-6p::gfp::ceh-6; odr-1::rfp]* | IS3122 |
| *gaIs245[col-34p::his-24::mcherry; unc-119(+)] V; oxIs12[unc-47p::gfp; lin-15(+)] X; fpEx788[egl-5p(1,3kb)::sox-2(antisens); rol-6(su1006)]* | IS3142 |
| *lin-40(ku285) V; oxIs12[unc-47p::gfp; lin-15(+)] X* | IS3146 |
| *sem-4(n1971) I; gaIs245[col-34p::his-24::mcherry; unc-119(+)] V; fpEx1062[let-413a::gfp::pest; myo-2p::gfp]* | IS3176 |
| *gaIs245[col-34p::his-24::mcherry; unc-119(+)] V; juIs8[unc-25p::gfp; lin-15(+)]* | IS3298 |
| *edIs6[unc-119p::gfp; rol-6(su1006)] IV; gaIs245[col-34p::his-24::mcherry; unc-119(+)] V* | IS3327 |
| *lin-17(n671)I; edIs6[unc-119p::gfp; rol-6(su1006)] IV; gaIs245[col-34p::his-24::mcherry; unc-119(+)] V* | IS3328 |
| *gaIs245[col-34p::his-24::mcherry; unc-119(+)] V; otIs118[unc-33p::gfp; unc-4(+)]* | IS3329 |

| | |
|---|---|
| *lin-17(n671)I; gaIs245[col-34p::his-24::mcherry; unc-119(+)] V; otIs118[unc-33p::gfp; unc-4(+)]* | **IS3330** |
| *lin-17(n671)I; gaIs245[col-34p::his-24::mcherry; unc-119(+)] V; juIs8 [unc-25p::gfp; lin-15(+)]* | **IS3335** |
| *jcIs1[ajm-1::gfp; rol-6(su1006)] IV; gaIs245[col-34p::his-24::mcherry; unc-119(+)] V* | **IS3339** |
| *lin-17(n671) I; fpIs110[lin-26p::gfp; rol-6(su1006)] IV; gaIs245[col-34p::his-24::mcherry; unc-119(+)] V* | **IS3349** |
| *lin-17(n671) I; jcIs1[ajm-1::gfp; rol-6(su1006)] IV; gaIs245[col-34p::his-24::mcherry; unc-119(+)] V* | **IS3357** |
| *gaIs245[col-34p::his-24::mcherry; unc-119(+)] V; fpEx1111[lim-6int4::gfp; coel::dsred]* | **IS3379** |
| *lin-17(n671) I; gaIs245[col-34p::his-24::mcherry; unc-119(+)] V; fpEx1062[let-413a::gfp::pest; myo-2p::gfp]* | **IS3383** |
| *lin-17(n671) I; gaIs245[col-34p::his-24::mcherry; unc-119(+)] V; fpEx1111[lim-6int4::gfp; coel::dsred]* | **IS3420** |
| *gaIs245[col-34p::his-24::mcherry; unc-119(+)] V; sox-2(syb737[gfp::linker::sox-2]) X* | **IS3423** |
| *wyIs75[unc-47p::dsred; exp-1p::gfp; odr-1p::rfp] III; vang-1(tm1422)X* | **IS3433** |
| *unc-73(e936) dpy-5(e61) I ; gaIs245[col-34p::his-24::mcherry; unc-119(+)] V ; oxIs12[unc-47p::gfp; lin-15(+)] X* | **IS3452** |
| *lin-17(n671) I; gaIs245[col-34p::his-24::mcherry; unc-119(+)] V; sox-2(syb737[gfp::linker::sox-2]) X* | **IS3457** |
| *sem-4(n1971) I; gaIs245[col-34p::his-24::mcherry; unc-119(+)] V; sox-2(syb737[gfp::linker::sox-2]) X* | **IS3458** |
| *dsh-1(ok1445) II; gaIs245[col-34p::his-24::mcherry; unc-119(+)] V; oxIs12[unc-47p::gfp; lin-15(+)] X* | **IS3464** |
| *egl-27(ok1670) II; wyIs75[unc-47p::dsred; exp-1p::gfp; odr-1p::rfp] III; him-5(e1490)V* | **IS3469** |
| *egl-5(n945) III; syIs50[cdh-3p::gfp; dpy-20(+)]* | **IS3475** |
| *egl-20(n585) IV ; gaIs245[col-34p::his-24::mcherry; unc-119(+)] V; oxIs12[unc-47p::gfp; lin-15(+)] X* | **IS3485** |
| *lin-44(n1792)I; gaIs245[col-34p::his-24::mcherry; unc-119(+)] V; oxIs12[unc-47p::gfp; lin-15(+)] X* | **IS3486** |
| *lin-44(n1792) I ; egl-20(n585) IV ; gaIs245[col-34p::his-24::mCherry; unc-119(+)] V ; oxIs12[unc-47p::gfp; lin-15(+)] X* | **IS3487** |
| *gaIs245[col-34p::his-24::mcherry; unc-119(+)] V; lim-6(nr2073) oxIs12[unc-47p::gfp; lin-15(+)] X* | **IS3490** |
| *par-1(zu310) gaIs245[col-34p::his-24::mcherry; unc-119(+)] V; oxIs12[unc-47p::gfp; lin-15(+)] X* | **IS3491** |
| *fmi-1(rh308) gaIs245[col-34p::his-24::mcherry; unc-119(+)] V; oxIs12[unc-47p::gfp; lin-15(+)] X* | **IS3511** |
| *dsh-1(ok1445) mig-5(tm2639) II; oxIs12[unc-47p::gfp; lin-15(+)] X* | **IS3512** |
| *wyIs75[unc-47p::dsred; exp-1p::gfp; odr-1p::rfp] III; gaIs245[col-34p::his-24::mcherry; unc-119(+)] V; sox-2(syb737[gfp::linker::sox-2]) X; fpEx1156[egl-5p(6.5kb)::nanobodyGFP::zif-1; coel::gfp; pBSK]* | **IS3521** |
| *gpr-1(ok2126) III; gaIs245[col-34p::his-24::mcherry; unc-119(+)] V; oxIs12[unc-47p::gfp; lin-15(+)] X* | **IS3530** |
| *sem-4(n1971)I; edIs6[unc-119p::gfp; rol-6(su1006)] IV; gaIs245[col-34p::his-24::mcherry; unc-119(+)] V* | **IS3537** |

| | |
|---|---|
| *sem-4(n1971)I; gaIs245[col-34p::his-24::mcherry; unc-119(+)] V; otIs118[unc-33p::gfp; unc-4(+)]* | **IS3539** |
| *gaIs245[col-34p::his-24::mcherry; unc-119(+)] V; ceh-6(syb972[gfp::linker::ceh-6]) X* | **IS3540** |
| *lin-17(n671) bxIs7[egl-5(6.5kb)::gfp; lin-15(+)] I; otIs173[rgef-1p::dsred2; ttx-3pB::gfp]III* | **IS3583** |
| *hT2[bli-4(e937) let-?(q782) qIs48] (I;III)/pop-1(q645)I ; gaIs245[col-34p::his-24::mcherry; unc-119(+)] V; oxIs12[unc-47p::gfp; lin-15(+)] X* | **IS3596** |
| *hT2[bli-4(e937) let-?(q782) qIs48] (I;III)/pop-1(q645)I; gaIs245[col-34p::his-24::mcherry; unc-119(+)] V; fpEx1111[lim-6int4::gfp; coel::dsred]* | **IS3600** |
| *otIs173[rgef-1p::dsred2; ttx-3pB::gfp]III; oxIs12[unc-47p::gfp; lin-15(+)] X* | **IS3604** |
| *lin-5(ev571)II; gaIs245[col-34p::his-24::mcherry; unc-119(+)] V; fpIs101[col-34p::ph::gfp; odr-1p::dsRed] X* | **IS3619** |
| *sem-4(n1971)I; gaIs245[col-34p::his-24::mcherry; unc-119(+)] V; lim-6(syb971[lim-6::linker::gfp])X* | **IS3632** |
| *lin-17(n671)I; gaIs245[col-34p::his-24::mcherry; unc-119(+)] V; lim-6(syb971[lim-6::linker::gfp])X* | **IS3669** |
| *gaIs245[col-34p::his-24::mcherry; unc-119(+)] V; lim-6(syb971[lim-6::linker::gfp])X* | **IS3677** |
| *lin-17(n671) I; gaIs245[col-34p::his-24::mcherry; unc-119(+)] V; ceh-6(syb972[gfp::linker::ceh-6]) X* | **IS3702** |
| *sys-1(q544) I; gaIs245[col-34p::his-24::mcherry; unc-119(+)] V; oxIs12[unc-47p::gfp; lin-15(+)] X* | **IS3718** |
